# High-Speed Low-Light *In Vivo* Two-Photon Voltage Imaging of Large Neuronal Populations

**DOI:** 10.1101/2021.12.07.471668

**Authors:** Jelena Platisa, Xin Ye, Allison M. Ahrens, Chang Liu, Ichun Anderson Chen, Ian G. Davison, Lei Tian, Vincent A. Pieribone, Jerry L. Chen

## Abstract

Monitoring spiking activity across large neuronal populations at behaviorally relevant timescales is critical for understanding neural circuit function. Unlike calcium imaging, voltage imaging requires kilohertz sampling rates which reduces fluorescence detection to near shot noise levels. High-photon flux excitation can overcome photon-limited shot noise but photo-bleaching and photo-damage restricts the number and duration of simultaneously imaged neurons. We investigated an alternative approach aimed at low two-photon flux, voltage imaging below the shot noise limit. This framework involved developing: a positive-going voltage indicator with improved spike detection (SpikeyGi); an ultra-fast two-photon microscope for kilohertz frame-rate imaging across a 0.4×0.4mm^2^ field of view, and; a self-supervised denoising algorithm (DeepVID) for inferring fluorescence from shot-noise limited signals. Through these combined advances, we achieved simultaneous high-speed, deep-tissue imaging of more than one hundred densely-labeled neurons over one hour in awake behaving mice. This demonstrates a scalable approach for voltage imaging across increasing neuronal populations.

## INTRODUCTION

To understand the nervous system, experiments are needed in which activity from large numbers of neurons can be measured in a detailed and comprehensive manner across multiple timescales during behavior. Current approaches to image neuronal activity at cellular resolution occupy two ends of a spectrum; recording large populations of neurons at very slow sampling rates (i.e. calcium imaging) or very high speed voltage imaging of small numbers of cells. However, calcium imaging is a poor proxy for action potential activity (Huang et al., 2021). To measure spiking accurately using voltage indicators, sampling must be at least an order of magnitude faster (∼400 Hz) (Sjulson and Miesenbock, 2007; Wilt et al., 2013) than calcium imaging (<15 Hz). To date, ultra-fast voltage imaging (1kHz) has only been achieved, simultaneously, in a small number (∼10) of neurons (Abdelfattah et al., 2019; Piatkevich et al., 2019; Villette et al., 2019; Wu et al., 2020). The goal of the present studies are to achieve high speed recording of a large number of neurons *in vivo*.

To optically resolve individual action potential in numerous neurons i*n vivo* faces two critical limitations. The first limit is determined by photon shot noise. Under the assumption of adequate indicator sensitivity, a sufficient number of genetically encoded voltage indicator (GEVI) molecules need to be excited for changes in fluorescence associated with action potential firing to be reliably detected above photon shot noise statistics. To achieve adequate signal-to-noise, current approaches for one-photon (1P) and two-photon (2P) voltage imaging have relied on “high photon flux” regimes in which excitation light is concentrated on a small population of tens of neurons (Adam et al., 2019; Villette et al., 2019). One-photon cellular resolution voltage imaging is achievable using widefield illumination and high frame-rate cameras but has limited depth penetration (Abdelfattah et al., 2019; Piatkevich et al., 2019). Due to light scattering, only neuronal populations at shallow depths or those sparsely labeled can be imaged with minimal signal degradation. Alternatively, 2P microscopy enables deeper imaging of densely labeled tissue at kilohertz acquisition rates but has been limited to a small field of views (FOVs) due to existing excitation and scanning strategies (Villette et al., 2019; Wu et al., 2020; Zhang et al., 2019).

High photon flux is partially driven by the characteristics of currently used 2P-compatible GEVIs that fluoresce at resting membrane potential and decrease their fluorescence during action potentials (negative slope fluorescence-voltage relationship, i.e., “negative going” indicator) (Chamberland et al., 2017; Jin et al., 2012; Villette et al., 2019). This means that, in addition to potentially mislocalized proteins, GEVI molecules at resting state also contribute to background fluorescence and light scatter (Abdelfattah et al., 2020; Piatkevich et al., 2019), further reducing detectable spike-related changes in fluorescence. Under both 1P and 2P imaging conditions, high photon flux prevents sustained voltage imaging due to rapid photobleaching which limits recording times to short durations (i.e., several minutes) (Abdelfattah et al., 2020; Piatkevich et al., 2019; Villette et al., 2019). Further, high photon flux excitation cannot be scaled for imaging larger neuronal populations. This is due to a second fundamental limit related to the total amount of excitation power that can be delivered into the brain without introducing photodamage (Podgorski and Ranganathan, 2016). Thus, when imaging at high speeds, the overall photon budget forces a trade-off in which the amount of excitation light available to each neuron decreases as the number of imaged neurons increases.

To achieve ultra-fast voltage imaging across large neuronal populations, these fundamental limits need to be overcome. We adopted a multidisciplinary approach to address this, integrating protein engineering, optical engineering, and deep learning. We developed a high-speed, positive-going 2P GEVI, a kilohertz-scanning, large-FOV 2P microscope, and a deep convolutional neuronal network for image denoising. Through this synergistic combination of technologies, we provide a new framework for population-level 2P voltage imaging in the awake behaving animal.

## RESULTS

### Development of positive-going two-photon compatible genetically encoded voltage indicator

We recently showed that we can manipulate the direction of the voltage-dependent fluorescence response of GEVIs by modifying amino acid residues determining the chromophore protonation state (Platisa et al., 2017) (**Figure 1A**). By mutating amino acid residues, D147A and H148A, within the green fluorescent protein (GFP) of the “negative-going” indicator ArcLight (Jin et al., 2012), we produced GEVI Marina (Platisa et al., 2017). This “positive-going” indicator is more advantageous for *in vivo* application due to lower background and improved photostability. However, despite optimal voltage sensitivity and optical properties, spike detection with Marina is limited by its suboptimal kinetics (tau ∼10 ms). Therefore, to develop a high-speed, positive-going voltage indicator, we devised a directed evolution strategy targeting the fastest available, negative-going, 2P compatible GEVIs, ASAP2f and ASAP3 (Chamberland et al., 2017; Villette et al., 2019).

**Figure 1.**
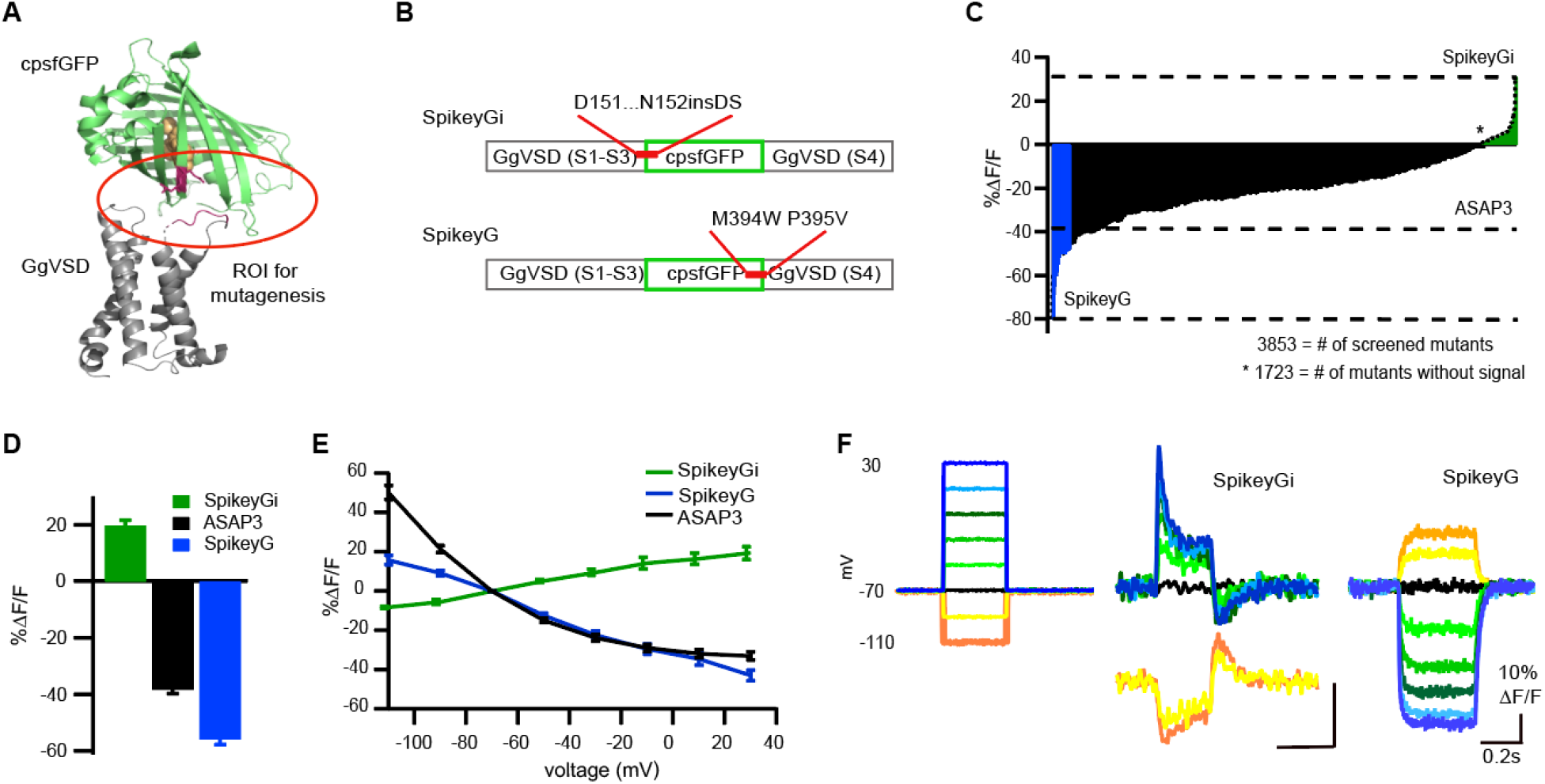
Design and functional characterization of green, dual-polarity, 2P compatible GEVIs. **(A)** A model of crystal structure for Spikey G and SpikeyGi based on crystal structures for cpEGFP (PDB 3EVP) and CiVSD (PDB 4G7Y). The region of interest (ROI) for targeted mutagenesis is labeled with the red circle. **(B)** A schematic showing insertion site for pair of amino acids DS between residues D151 and N152 within FP that produced SpikeyGi (upper), and double point mutation M394W P394V within VSD that produced SpikeyG. **(C)** Distribution of fluorescence responses of ASAP3 mutants (in black) in response to field simulation-based functional screening in expressing electrically active HEK293 cells. Positive-going SpikeyGi variants are shown in green, negative-going SpikeyG variants are shown in blue, and parent ASAP3 in black. **(D)** The amplitude of fluorescence response to 300ms 100mV depolarization voltage step in transiently expressing HEK293 cells of novel GEVIs, SpikeyGi (*n* = 8 cells) and SpikeyG (*n* = 5 cells) vs. parent indicator ASAP3 (*n* = 8 cells). The color scheme is the same as in [C]. Data are plotted as mean ± SEM. **(E)** V-F curve showing fluorescence response of novel indicators across a range of voltage steps for SpikeyGi (*n* = 3 cells), SpikeyG (*n* = 5 cells), and parent indicator ASAP3 (*n* = 5 cells). For all cells, 300ms voltage steps of −40 to +100 mV were applied in increments of 20 mV from a resting potential of −70 mV. The color scheme is the same as in [C]. Data are plotted as mean ± s.e.m. **(F)** Example of traces showing fluorescence response to series of voltage steps recorded from HEK293 transiently expressing novel indicators, SpikeyGi and SpikeyG.

We began by creating site-directed mutagenic libraries of amino acid residues within the ASAP2f fluorescent protein (S149, and H150; from the starting Met of ASAP2f) that are homologous to those in ArcLight/Marina (D147 and H148; numbering from starting Met of SuperEcliptic pHluroin GFP). Additional libraries targeted four residues at the linker region between the voltage-sensitive and FP domains (L145, S146, F147, and N148). Functional screening showed that most variants produced signals that were either smaller or similar to the parent constructs. However, as seen with Arclight and Marina, mutation of H150 residue produced several variants with reversed signal polarity, i.e., a modest increase in fluorescence upon depolarization (data not shown). We then used primers with the degenerative codons NNKNNK to create an insertional library targeted between residues H151 and N152 in ASAP3 (**Figure 1B**). To ensure that all 400 potential variants were screened, we tested 1104 mutants produced in two separate PCR reactions. Out of all mutants screened, a ∼150 positive-going variants were detected (**Figure 1C**). In one mutant, a large ΔF/F_0_ of +30% was observed using field stimulation (**Figure 1C-D**). Subsequently, simultaneous high-speed 1P imaging (500-1000Hz) and whole-cell patch-clamp electrophysiology of HEK cells transiently expressing this mutant showed an average +18.7 ± 1.1% ΔF/F_0_ response to 100 mV step depolarization (*n* = 8 cells) (**Figure 1E-F**). Sequence analysis of this indicator revealed the insertion of amino acids DS; this indicator was named SpikeyGi.

Along with our efforts to generate positive-going 2P GEVIs, we developed additional negative-going 2P GEVIs with improved signal characteristics. We created site-directed libraries targeted to the 26 amino acid residues between the circularly permuted sfGFP and S4 sequences of the voltage-sensitive domain of ASAP3 (residues 370-395 in ASAP3 from starting Met, **Figure 1A-B**). Two subsequent mutagenesis and functional screening rounds (at positions 394 and395) led us to the double mutant M394W P395V that shows sensitivity optimized to detect depolarizations **(Figure 1E and F)**. Compared to parent GEVI ASAP3, this novel variant, named SpikeyG, showed increased sensitivity to transient 100mV step depolarizations (ΔF/F_0_: SpikeyG, -55.7 ± 1.9%, *n* = 6 cells; ASAP3, -38.1 ± 1.6%, *n* = 8 cells, Student’s *t*-test; *p*<0.00002; **Figure 1D**). At the same time, sensitivity to transient -50mV step hyperpolarization (−110mV voltage step from -70mV holding potential) decreased for SpikeyG (ΔF/F_0_: SpikeyG, 24.6 ± 2.1%, *n* = 5 cells; ASAP3, 51.4 ± 4.2%, n = 8 cells. Student’s *t*-test; *p*< 0.0002; **Figure 1E**). Overall, our directed mutagenesis of fast two-photon-compatible GEVIs produced two improved candidates with the potential for increased sensitivity for spike detection.

### *In vitro* characterization of SpikeyG and SpikeyGi with high photon flux excitation

To determine the suitability of SpikeyG or SpikeyGi for action potential detection in neurons, we first characterized responses of SpikeyG and SpikeyGi *in vitro* with simultaneous whole-cell patch clamp electrophysiology and two-photon imaging. We targeted layer 2/3 neurons in brain slices from animals virally expressing either SpikeyG or SpikeyGi. Fluorescence responses across different membrane potentials were measured under voltage-clamp mode. We imaged single neurons under high photon flux conditions using a conventional two-photon microscope with an 80 MHz repetition rate laser source, similar to previous studies (Bando et al., 2019; Chamberland et al., 2017). Using frame scanning (128×85 pixels, 0.3-0.6ʼμm/pixel. 2.8ʼμs dwell time, 35-45mW average power), the targeted cell received ∼5×10^5^ pulses per 50ms time bin (**Figure 2A**). SpikeyG showed linear changes in response to voltage steps, with negative steps producing an increase in fluorescence and positive steps producing a decrease in fluorescence. SpikeyGi responded in the opposite direction with positive steps increasing fluorescence (**Figure 2B-C**). At +50mV (−120mV voltage step from -70mV holding potential), SpikeyGi showed larger magnitude fluorescence changes than SpikeyG (ΔF/F_0_: SpikeyGi, 29.4 ± 4.7%; SpikeyG, -14.6 ± 5.1%). We next characterized single action potential responses of SpikeyG and SpikeyGi triggered at 100-ms intervals (average of 10 traces). Line scans at 3-5 kHz were performed along the cell membrane (24-54 pixels, 2.8 ʼs dwell time), equivalent to ∼9000 excitation pulses per 1ms time bin. Both GEVIs showed clear responses to action potentials with similar kinetics. SpikeyGi exhibited greater magnitude peak responses compared to SpikeyG (ΔF/F_0_: SpikeyGi, 32.6 ± 0.9%; SpikeyG, -21.2 ± 0.4%; Student’s *t*-test, *p* < .001) (**Figure 2D**). These results demonstrate that both SpikeyG and SpikeyGi are capable of reporting single APs under high photon flux conditions.

**Figure 2.**
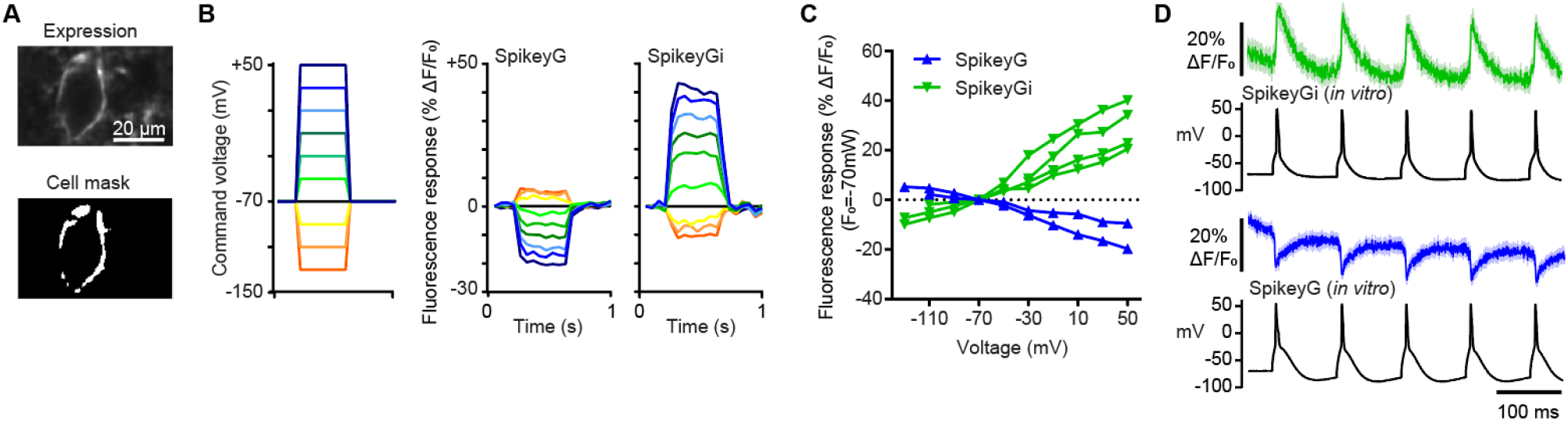
*In vitro* performance of SpikeyGi and SpikeyG. **(A)** Example of field of view of neuron imaged for slice experiments (top). ROI mask used for image analysis (bottom). **(B)** Example command voltages applied during voltage clamp mode (left). Corresponding fluorescence responses measured in example SpikeyG and SpikeyGi cells). **(C)** Fluorescent responses to steady-state voltage steps in slice electrophysiology for individual cells (Normalized to -70 mV; SpikeyG, *n*=2 cells; SpikeyGi, *n*=4 cells; 10 trials per step). **(D)** Fluorescent responses to 10Hz action potential trains evoked by current injection *in vitro (n =* 10 trials). Shaded region; s.e.m.

### Ultra-fast two-photon microscope design and performance

To perform two-photon voltage imaging across a large population of neurons, we sought to design an ultra-fast two-photon microscope capable of imaging a 400×400ʼμm^2^ field-of-view (FOV) at a kilohertz frame rate. While existing ultra-fast two-photon microscopes operate in a high-photon photon flux regime, we set out to construct a system optimized for “low-photon flux” excitation (**Figure 3A**). Two-photon excitation is achieved through pulsed lasers. Hence, the pulse repetition rate determines the total FOV that can be excited within a one-millisecond time bin. A minimum of one pulse is needed per imaging voxel to cover the entire FOV. To increase the FOV size while maintaining full coverage, the effective repetition rate of the imaging system needs to increase proportionally. This can be achieved through either temporal or spatial multiplexing. Temporal multiplexing creates multiple excitation beamlets that are delayed in time such that the resulting fluorescence detected by a single photomultiplier tube (PMT) can be disambiguated by their timing (Amir et al., 2007; Chen et al., 2016; Cheng et al., 2011; Clough et al., 2021). However, the degree of temporal multiplexing is limited by the fluorescence lifetime of the excited fluorophore which places an upper limit on the effective repetition rate using this approach. In contrast, there is no limit to the effective repetition rate formed by spatial multiplexed beamlets in which multiple pulses are delivered into different regions of the tissue simultaneously (Kim et al., 2007; Zhang et al., 2019). Resolution of spatially multiplexed beamlets does require spatial detection using cameras or multi-anode PMTs (MAPMTs). Consequently, spatial multiplexing is depth limited as crosstalk between neighboring detectors increases with depth due to scattered fluorescence. This spatial crosstalk can be reduced by increasing the spacing of the beamlets at the sample.

**Figure 3.**
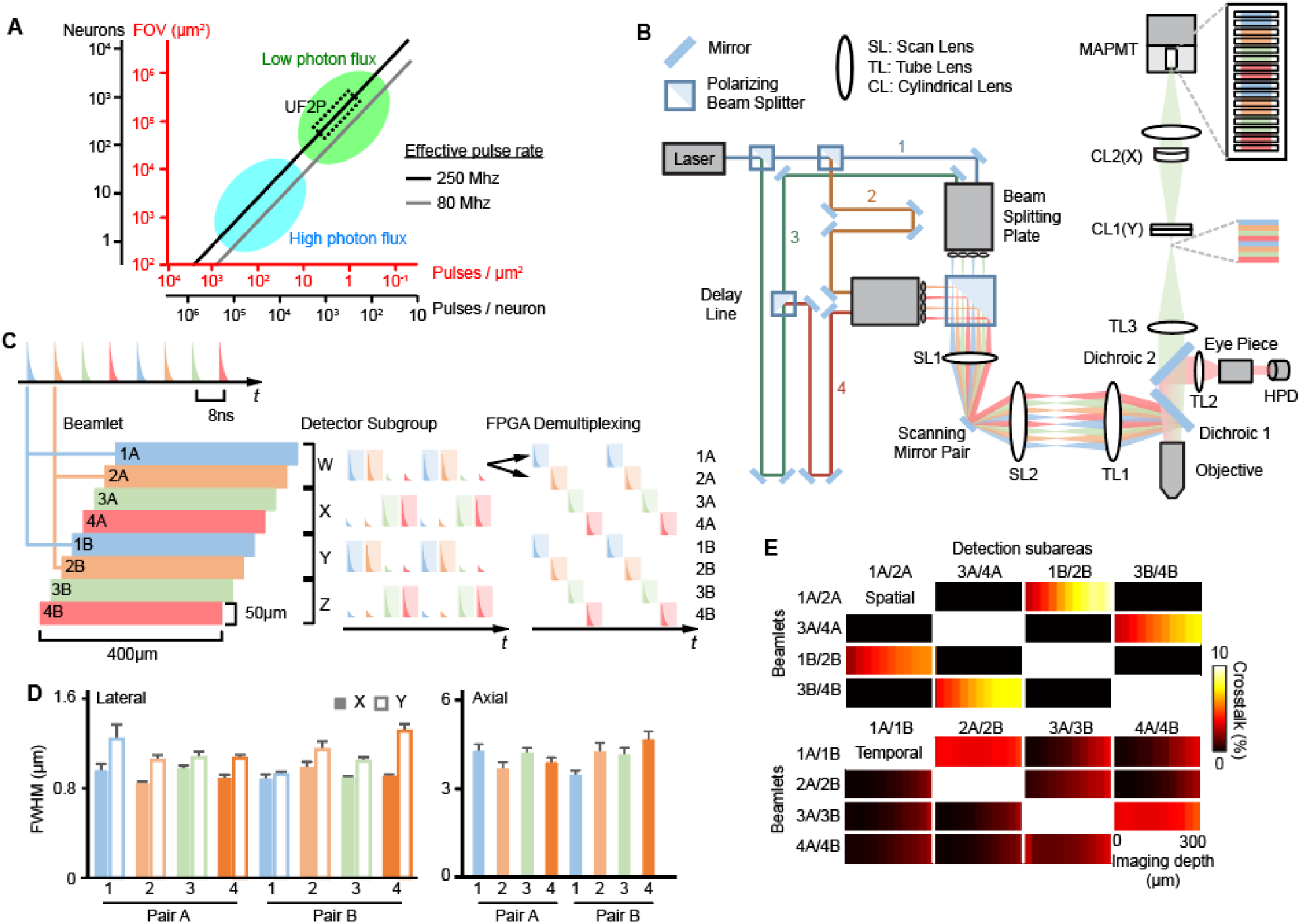
Design and performance of ultra-fast two-photon microscope. **(A)** Relationship between excitation pulses per neuron and total neurons imaged at 1 kHz sampling rate assuming 1 ʼm^2^ voxel across different effective pulse rates. **(B)** Schematic of the ultra-fast two-photon microscope. Laser beam was first split into 4 beamlets (blue - 1, orange - 2, green - 3, red - 4) using polarizing beam splitters (PBS). Beamlets were temporally multiplexed using delay lines and then split into spatially multiplexed beamlet pairs. Using two beam splitting plates, beamlets were spatially arranged, combined with a 2-inch PBS into a linear arrangement, and projected onto the objective back pupil. Scan and tube lenses were matched to the beam diameter between resonant/galvo scanner pair and the size of the objective back aperture. The detection path enabled single color imaging with all 8 beamlets or dual color imaging with a single beam. A dichroic separated green from red fluorescence. Red fluorescence excited from a single beam was detected with a single hybrid PMT. For green fluorescence, cylindrical lenses in the detection path reshaped the collected fluorescence to match a linearly arranged 16 × 1 MAPMT detector. Signals from each anode were independently collected when imaging using 8 beamlets or summed when excited with a single beam. **(C)** Schematic of detection and demultiplexing algorithm. The 16 anodes on the MAPMT were summed into 4 detector subgroups (W, X, Y and Z). Each detector subgroup received photons from 2 temporally multiplexed subareas that were subsequently demultiplexed using FPGA programmed digital gates (shaded area). Additional time-dependent gating was implemented on the FPGA to minimize spatial multiplexed crosstalk from neighboring detector subgroups. (**D**) Lateral and axial PSF measurements for each beamlet. (*n*=7-11 beads per beamlet). Error bar indicates the s.e.m. (**E**) Average detected crosstalk as a function of imaging depth due to spatial multiplexing (top panel) or temporal multiplexing (bottom panel). (*n* =5 imaging stacks). See also **Figure S1-S2**.

To maximize the effective repetition rate used in the UF2P microscope, spatial and temporal multiplexing were combined in the same system (**Figure 3B, S1**). For the excitation source, we selected a 920nm 31.25MHz fiber laser which enables beamlets to be temporally multiplexed four times at 8-ns pulse intervals, sufficient to resolve GFP-based indicators with minimal cross talk. Compared to 80 MHz Ti:sapphire lasers traditionally used for two-photon imaging, the lower repetition rate also provides >2.5x greater pulse energy at the same average power, providing more efficient excitation per laser pulse (Charan et al., 2018). Each temporally multiplexed beamlet was then split into a pair of spatially multiplexed beams positioned 200μm apart at the sample to minimize scatter-related crosstalk. This generated a total of 8 beamlets (4 temporal X 2 spatial). The result is an illumination source with an effective 250 MHz repetition rate, which is >3x higher than traditional systems scanning a single beam with 80MHz repetition rate. For raster scanning, we used a resonant mirror with 24kHz line rates for x-scanning and a galvanometric mirror for y-scanning. Since fast scanning is achieved along the x-axis, the beamlets were linearly arranged along the slower y-axis. The four temporally multiplexed beamlet pairs were spaced 50μm apart at the sample (**Figure 3C**). Thus, each beamlet scanned a 400×50 μm sub-area, when tiled together, resulting in a total FOV of 400×400μm^2^. Each sub-area was slightly offset in the x-axis as a result of the patterning of the beamlets with respect to the position of the resonance scanner. By scanning 24 lines per subarea either, unidirectionally or bidirectionally, we achieved 803 or 1000 Hz frame rates, respectively. A linearly arranged MAPMT and matching detection optics were designed to project the imaging plane onto the detectors such that each anode collected fluorescence from a corresponding spatially multiplexed beamlet pair. The fluorescence collected from the MAPMT was subsequently demultiplexed to resolve the temporally multiplexed beamlets while minimizing crosstalk fluorescence in neighboring anodes.

We first evaluated the optical performance of the microscope. To check if each beamlet provides similar excitation and resolution in each sub-area, the point spread function (PSF) of each beamlet was individually measured using fluorescent beads (*n*=7-11 beads per beam, **Figure 3D, S2**). The optical performance was similar for each sub-area. Across all sub-areas, the microscope achieved an average PSF of 0.9±0.1μm(X)/1.1±0.1μm(Y) lateral resolution and 4.1±0.4μm axial resolution, demonstrating sub-cellular resolution performance. To determine the degree of temporal and spatial cross talk across all 8 beamlets in scattering tissue, a cranial window was implanted in a mouse with virally expressed SpikeyGi in the primary somatosensory cortex (S1). SpikeyGi fluorescence was measured as a function of cortical depth for each beam across all detected sub-areas (**Figure 3E**). We observed that crosstalk caused by spatial multiplexing increased as a function of imaging depth but remained under 10% as far as 300μm below the pial surface. Crosstalk as a result of temporal multiplexing remained under 5%, independent of imaging depth. Overall, this demonstrates that the UF2P microscope can achieve large FOV kilohertz frame scan imaging into deep tissue.

### Self-supervised denoising improves action potential detection below photon shot noise limits

While increasing the effective pulse rate with the UF2P microscope enables imaging FOVs to be increased while maintaining high frame rate, photon flux is still magnitudes lower then high photon flux regimes. Assuming a photon flux of ∼0.1 ʼs dwell time per μm^2^ voxel, each neuron only receives ∼200 excitation pulses per 1ms time bin. Under such imaging conditions, shot noise dominates pixel-wise measurements. Recently, self-supervised deep learning denoising algorithms have been developed to remove independent noise sources in calcium imaging data without any ground-truth “clean” (high SNR) measurements (Lecoq et al., 2021; Li et al., 2021). We expanded upon this approach, developing a deep convolutional neural network (CNN) to denoise voltage imaging data (DeepVID) (**Figure 4A**). DeepVID combines self-supervised denoising frameworks that infers the underlying fluorescence signal based on a learned model of the independent temporal and spatial statistics of the PMT measurements that is attributable to shot noise (Krull et al., 2018; Lecoq et al., 2021). The CNN was trained to estimate a single center frame (*N*_*0*_) based on information from prior (*N*_pre_) and subsequent (*N*_post_) neighboring frames within a time series. Simultaneously, the CNN was trained to estimate a few “blind” pixels (*p*_blind_, in %) within the central frame based on information from all remaining pixels within that frame. Based on the rise time of SpikeyG and SpikeGi and imaging frame rate of the UF2P microscope, a model with *N*_pre_=3 and *N*_*post*_=3 was chosen to maximize inference accuracy while preserving the fast action potential kinetics of the indicators. We first assessed the frame-to-frame variability in fluorescence signal in the raw data and confirmed that the fluctuations in each pixel is proportional to the square root of the mean fluorescence (**Figure 4B**), as expected for shot noise limited signals. DeepVID drastically reduced the frame-to-frame variability, resulting in a 15-fold improvement in SNR when comparing denoised and raw image data (SNR: 0.567 ± 0.002, raw; 8.858 ± 0.027, denoised, n = 8,000 pixels) (**Figure 4C**). By breaking this fundamental noise constraint, the underlying fluorescence signal can be more accurately inferred at individual time points (**Figure 4D**).

**Figure 4.**
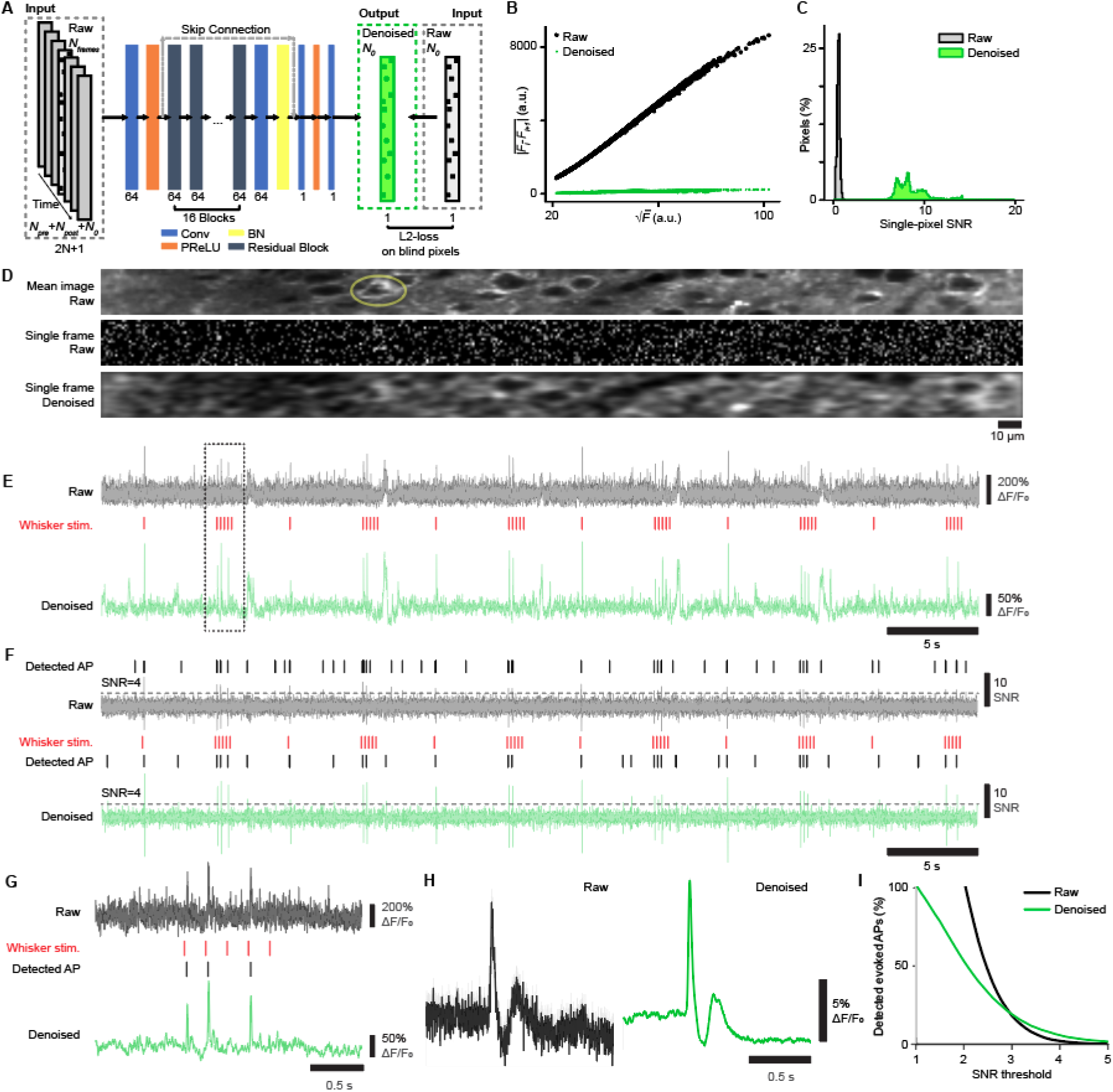
DeepVID reduces photon shot noise to improve action potential detection. **(A)** Training strategy and network structure of DeepVID. The voltage signals from blind-spot pixels in the central frame (*N*_*0*_) are inferred from the rest of unmasked pixels in the central frame and the neighboring frames (*N*_*pre*_ and *N*_*post*_) in the input image data. DeepVID is a deep convolutional neural network with residual blocks, where Conv is 2D convolution, BN is batch normalization, and PReLU is the parametric rectified linear unit. (B) Frame-to-frame noise (mean of the absolute intensity difference) as a function of fluorescence signal (square-root of the mean intensity) for raw vs. denoised pixels for representative image data. (C) Distribution of pixel-level SNR (temporal mean over S.D. for each pixel time series) from a representative raw and denoised data. **(D)** Example of single frame image denoising with DeepVID. Images from one subarea acquired with the UF2P microscope. Raw *in vivo* image averaged across 1000 frames (top panel) showing SpikeyGi-expressing neurons. Single frame is shown before (middle panel) and after denoising with DeepVID (bottom panel). **(E)** Raw and denoised fluorescence traces from neuron [circled in D]. Air puff whisker stimulus are shown. (**F**) Putative spike events based on SNR levels in raw and denoised traces. Detected events at SNR>4 are shown. **(G)** High temporal resolution view of example raw and denoised traces [box in E] showing spike-related fluorescence changes. **(H)** Average raw and denoised fluorescence traces in response to single air puff stimulus. **(I)** *In vivo* detection of sensory-evoked APs with SpikeyGi across SNR thresholds for raw and denoised traces (*n* = 214 cells, 3 animals). Shaded region in E, F,G and H equals S.E.M.

To assess how DeepVID improves the reliability of spike detection in voltage imaging data, we measured neuronal responses using SpikeyGi in S1 during whisker stimulation. Whisker deflections using air puffs produce well-timed single AP responses in L2/3 S1 neurons (Feldmeyer et al., 2012). Whisker stimulation was delivered in the form of single or trains of 5 air puffs to the contralateral whisker pad. We compared raw and denoised fluorescence traces. The reduction in shot noise fluctuations in denoised traces readily allowed for the identification of potential sensory-evoked and non-evoked spiking events (**Figure 4E**). Fluorescence traces were converted into timeseries of SNR levels used for spike detection (**Figure 4F**). Denoising appeared to improve the SNR of putative spike events. Analysis of the fluorescence response (spiking and non-spiking) to single air puffs in denoised vs. raw traces show that peak responses were increased in denoised traces, owing to better estimates of the baseline fluorescence levels (ΔF/F_0_ raw: 9.3 ± 1.0%; denoised, 11.1 ± 0.2%; Student’s t-test, p < 0.001) (**Figure 4G-H**). We compared the percent of putative sensory-evoked spikes detected at varying SNR thresholds (**Figure 4I**). At low SNR levels (< 3), the percent of detected spikes was highly overestimated in raw traces whereas the likelihood of false positives was greatly reduced in denoised traces. In contrast, denoising improved the detection of spikes at high SNR thresholds (> 3) compared to raw traces. Overall, these results demonstrate that reduction in shot noise provided by DeepVID significantly improves the reliability for spike detection in 2P voltage imaging data.

### Positive-going GEVIs outperforms negative-going GEVIs *in vivo*

Using DeepVID, we next compared responses of SpikeyG and SpikeyGi *in vivo* under low photon flux conditions. Imaging across the full 400 × 400 μm^2^ FOV, cells were imaged at similar laser power compared to *in vitro* conditions (∼30 mW per beamlet) but at lower resolution (x: 1.0 μm/pixel, y: 2.1 μm/pixel). We first compared the fluorescent responses to single air puffs (**Figure 5A**). SpikeyGi showed larger peak responses compared to SpikeyG (ΔF/F_0_: SpikeyGi, 11.1 ± 0.2%; SpikeyG, 5.6 ± 0.1%; Student’s t-test, p < 0.001). For SpikeyGi, the detection of APs across different SNR thresholds were consistently higher than SpikeyG repeated measures ANOVA, group interaction: F = 407.41, *p* < 0.001) (**Figure 5B**). We further evaluated the ability to perform population imaging with SpikeyGi using the UF2P microscope. We simultaneously imaged 129 SpikeyGi neurons across the FOV and could identify sensory-evoked APs (SNR > 4) across neurons to both single and trains of air puffs (**Figure 5C-D, S3**). We evaluated the temporal fidelity of SpikeyGi responses to 5-10 Hz trains of the air puff. Since spiking probability to whisker stimulation can be variable from cell-to-cell in S1 neurons, we generated average traces to detected APs for each air puff in the stimulus train (**Figure 5E**). At both 5 and 10 Hz stimulation frequencies, the averaged traces show well isolated spike responses to each air puff within the stimulus train. These results demonstrate that SpikeyGi in combination with the UF2P microscope is suitable for population-level voltage imaging.

**Figure 5.**
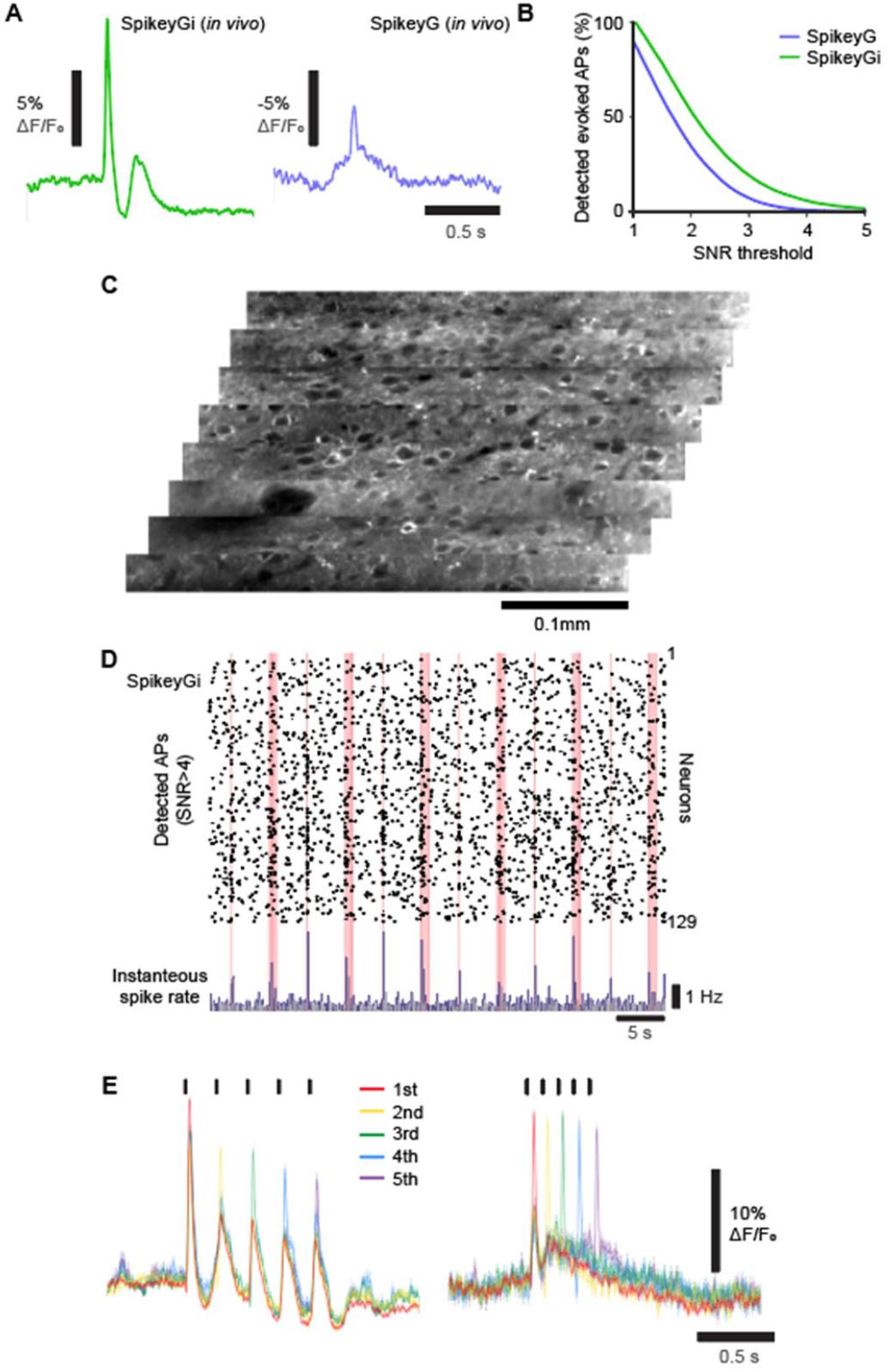
SpikeyGi outperforms SpikeyG for *in vivo* two-photon population imaging. **(A)** Average raw and denoised fluorescence traces in response to single air puff stimulus for SpikeyGi and SpikeyG. (*n* = 214 cells, 3 animals; SpikeyGi, 135 cells, 2 animals; SpikeyG). **(B)** *In vivo* detection of sensory-evoked APs across SNR thresholds for SpikeyGi and SpikeyG. **(C)** Example FOV from UF2P microscope of L2/3 neurons expressing SpikeyGi. **(D)** Detected *in vivo* spike trains at SNR>4 for 129 simultaneously imaged neurons expressing SpikeyGi. Red lines indicate air puffs. Instantaneous spike rate across the population are shown at the bottom. **(E)** Average fluorescence responses to detected APs (SNR >4) for individual air puffs in 5Hz (left) or 10Hz (right) stimulus trains (*n* = 214 cells, 3 animals, 5Hz stimuli, and *n* = 206 cells, 3 animals, 10Hz stimuli). Shaded region; s.e.m. See also **Figure S3**.

### Low photon flux enables sustained population-level voltage imaging

We finally assessed the capacity to perform sustained *in vivo* voltage imaging at low photon flux using the UF2P microscope. We first tested photobleaching seen SpikeyGi and SpikeyG under *in vitro* high photon flux conditions. This was performed in a 512 × 512-pixel field of view at 1 Hz with laser power of 35mW. Intermittent imaging (10s on, 5s off) was performed across 12.5 minutes. Fluorescence was normalized to the first frame of the recording. SpikeyGi showed little reduction in fluorescent output over the course of the recording and showed significantly less bleaching than SpikeyG (repeated measures ANOVA group x time interaction: *F*_749, 14231_ = 2.221, *p* < 0.0001) (**Figure 6A**). Next, we compared photobleaching rates *in vivo* at low photon flux conditions using UF2P microscope under similar intermittent imaging conditions (9s on, 4s off) across 60 min. For both SpikeyGi and SpikeyG, the fluorescence rapidly decreased by 11-13% within the first 5 minutes but then more slowly reduced to 22-25% after one hour. Unlike under high photon flux conditions, no difference in photobleaching was observed between SpikeyGi and SpikeyG under low photon flux conditions. These results demonstrate that low photon flux excitation benefits both positive- and negative-going indicators. For SpikeyGi, we assessed the action potential (AP) detection rate to air puff stimulation across the one hour of imaging (**Figure 6B-C**). Despite the changes in fluorescence, no differences in action potential detection were observed across the period.

**Figure 6.**
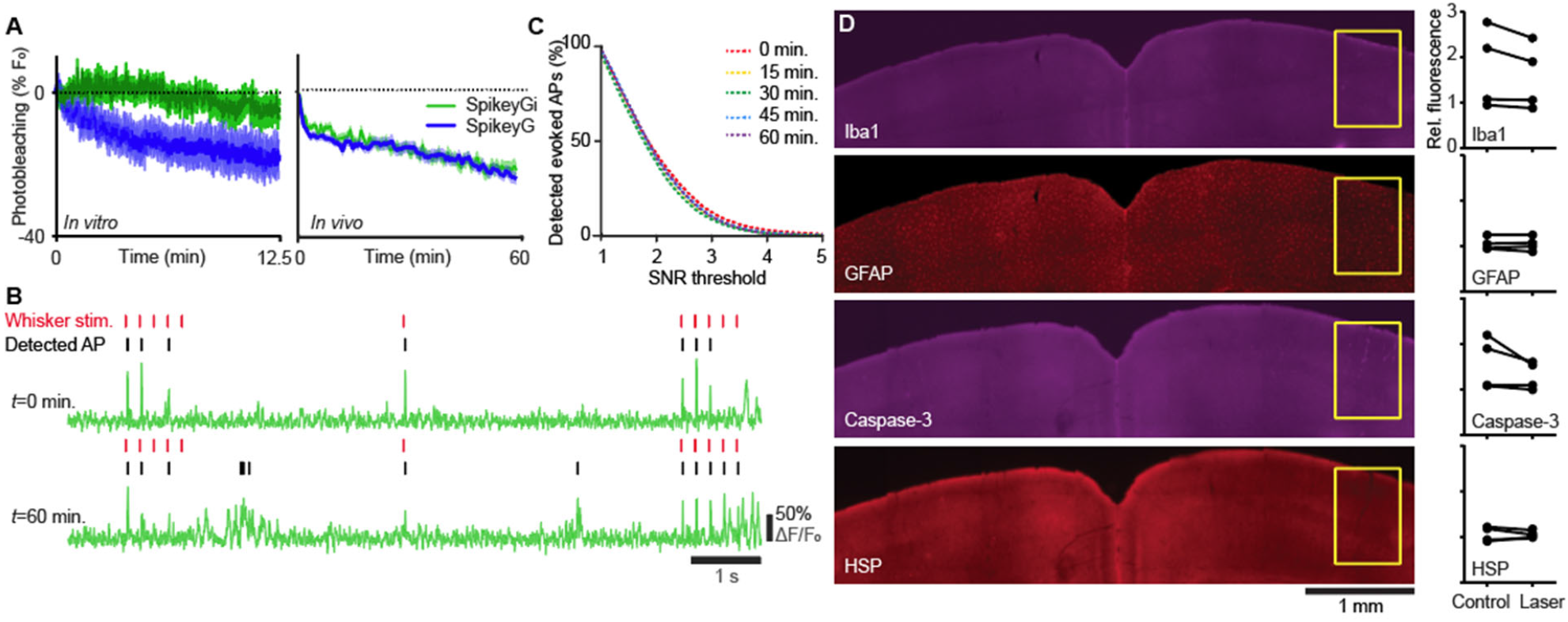
Low photon flux excitation facilitates sustained two-photon voltage imaging. **(A)** Left panel shows *in vitro* photobleaching curves for SpikeyGi and SpikeyG under high photon flux conditions (*n* = 8 cells, 1 FOV, SpikeyGi; 13 cells, 3 FOV, SpikeyG). Right panel *In vivo* photobleaching curves for SpikeyGi and SpikeyG under low photon flux conditions (*n* = 33 cells, 3 FOV, SpikeyGi; 33 cells, 3 FOV, SpikeyG). **(B)** Example SpikeyGi fluorescence traces and detected action potentials (SNR>4) of a neuron across 1 hour of intermittent *in vivo* imaging (9s on, 4s off). **(C)** *In vivo* detection of sensory-evoked APs across SNR thresholds for SpikeyGi across one hour of imaging. **(D**) Left panels show example coronal sections of immunostained tissue assaying photodamage after one hour of sustained imaging. Yellow box denotes imaged region. Right panels show relative fluorescence in imaged region compared corresponding to contralateral region areas across immunostained tissue (*n* = 4 animals). Scale bar: 1 mm. Shaded region; s.e.m.

Given that the UF2P microscope delivers a total of 240mW across 8 beams (30mW per beamlet) during *in vivo* imaging, we tested for signs of photodamage or toxicity after 1 hour of sustained imaging (Podgorski and Ranganathan, 2016). Animals were perfused 16 hours after imaging, and immunostaining was performed for four markers of tissue damage and inflammation: astrocytic (anti-GFAP), microglial (anti-Iba1), heat shock (anti HSP-70/72), and apoptotic pathway activation (anti-Caspase-3). We measured average fluorescence levels in the laser-exposed areas of the cortex and in control areas of the cortex which were not exposed to laser scanning (**Figure 6D**). We found that laser exposure did not produce an increase in any of the four markers of photodamage. There was no significant difference between laser-treated and control areas of the cortex for all four markers (paired Student’s *t*-test, *n*=4: GFAP, t = 2.27, *p* = 0.11; Iba1, t = 0.06, *p* = 0.95; HSP, t = 1.75, *p* = 0.18; Caspase-3, t = 0.39, *p* = 0.72). In summary, low photon flux imaging using UF2P microscope in combination with SpikeyGi provides safe and sustained population-level voltage imaging.

## DISCUSSION

Here we developed a novel, multidiscipline approach to chronically record fast voltage transients from large sets of neurons in deep brain tissue in behaving animals. The performance gains of the system are possible by the combination of three independent novel advances; a high performing, positive-going, fast, voltage indicator (SpikeyGi), an ultra-fast, large FOV multiphoton microscope and a purpose-built AI denoising process (DeepVID). With these innovations we demonstrate sustained large-scale, ultra-fast, two-photon, voltage imaging for the first time.

Our approach uses a positive fluorescence-voltage slope relationship GEVI (SpikeyGi). During its characterization *in vitro* under high photon flux illumination, Spikey Gi and its “negative going” counterpart (SpikeyG) showed similar SNR performance. However, under the low photon flux imaging conditions of our UF2P microscope, SpikeyGi significantly outperforms its “negative-going” counterpart. While we have not yet pursued the basis for this difference, we hypothesize reduced baseline noise and the higher photon count during action potentials combined in SpikeGi result in significantly improved SNR. This indicates that *in vitro* characterization is not the best predictor of *in vivo* performance, but rather an approach which combines probe and imaging method engineering is likely to be more successful.

In the study, we utilize temporal and spatial multiplexing to develop the UF2P microscope. By increasing the effective pulse rate of the excitation source while maintaining low detection cross talk, the UF2P microscope can achieve kilohertz frame scanning in deep tissue at twice the FOV of current kilohertz-scanning two-photon systems (**Table 1**). The FOV size and imaging depth are comparable to standard 2P microscopes performing calcium imaging at very low speed. This inherently results in lower photon flux per imaged neuron. Low photon flux excitation results in reduced photobleaching and photodamage, enabling sustained and chronic voltage imaging. However, it is challenging to achieve reliable fluorescence imaging under shot noise limiting conditions.

**Table 1.**
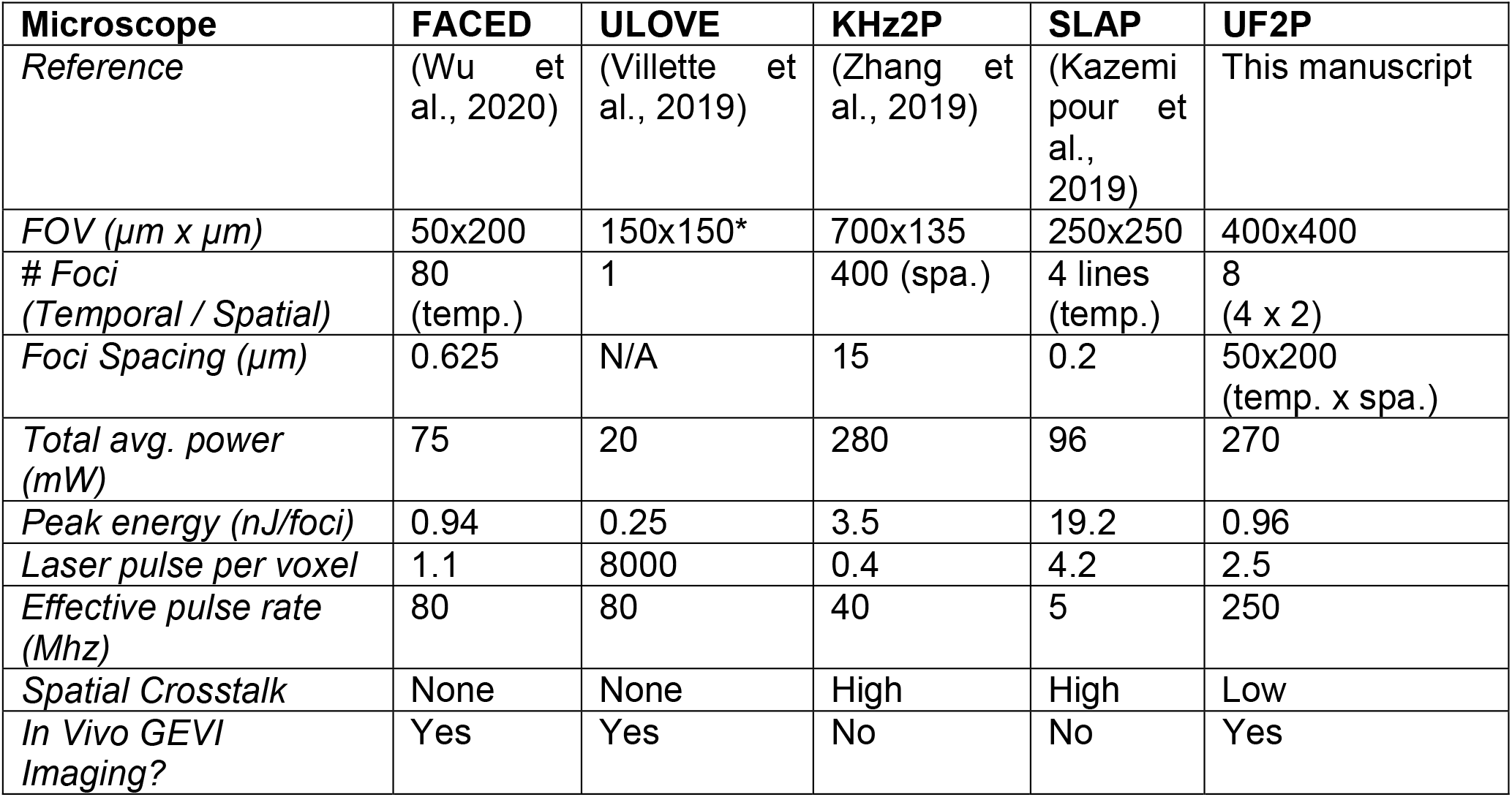
Comparison of kilohertz-scanning two-photon microscopes. *indicates Random access scanning.

To improve imaging at low photon flux levels we developed a self-supervised, deep learning denoising method (DeepVID). DeepVID denoising was achieved without sacrificing spatial or temporal resolution and without access to “ground-truth” high-SNR measurements. It achieved a 15-fold increase in the single-pixel SNR, which is comparable to the improvements reported in 2P calcium imaging (Lecoq et al., 2021; Li et al., 2021). Applying DeepVID to sensory-evoked measurements revealed more reliable spike detections.

The combination of these tools represents a new direction of further technology development to scale up two-photon voltage imaging. For the UF2P microscope, FOV can be further increased while maintaining overall photon budget levels by a combination of additional judiciously placed spatially multiplexed beamlets to increase the effective repetition rate and beam shaping to improve excitation efficiency (Demas et al., 2021; Weisenburger et al., 2019). Ongoing developments in denoising can further reduce the photon flux needed for reliable detection. The denoising performance of DeepVID can be improved by incorporating more advanced network architectures, such as spatiotemporal convolution and attention module (Vaswani et al., 2017). Improvements in SpikeyGi can further reduce photon flux requirements by increasing fluorescence response amplitude to membrane potential changes, decreasing resting fluorescence to reduce background, and improving subcellular localization to reduce inactive SpikeyGi molecules. Overall, low-photon flux imaging pushes the boundaries for monitoring neuronal activity across spatial and temporal scales. Additionally, this framework can be adopted for other *in vivo* imaging applications that track signal changes in molecules of low abundance across large areas at fast temporal resolutions.

## ACKNOWLEDGEMENTS

We thank K Khait for assistance in Scope software development; N Manjrekar for assistance in photodamage experiments; Pieribone Laboratory scientific staff J. Wojciekofsky, L. Delgado, and P. O’Brien for technical assistance; the Pierce Laboratory Instrument shop including J. Buckley, A. Wilkins, T. D’Alessandro, and A. DiRubba for help with instrumentation. This work was supported by grants from a NARSAD Young Investigator Grant from the Brain & Behavior Research Foundation (J.L.C.), the Richard and Susan Smith Family Foundation (J.L.C.), Elizabeth and Stuart Pratt Career Development Award (J.L.C.), the Whitehall Foundation (J.L.C.), NSF Neuronex Neurotechnology Hub NEMONIC #1707287 (J.L.C.), NIH New Innovator Award DP2NS111134 (J.L.C.), NIH BRAIN Initiative Awards R01NS109965 (J.L.C.), UF1NS107705 (J.L.C. and V.A.P.), R21EY030016 (L.T.), U01NS103517 (V.A.P), U01NS090565 (V.A.P), and DARPA N6600117C4012 (V.A.P.), and N660119C4020 (V.A.P.).

## AUTHOR CONTRIBUTIONS

J.L.C. and V.A.P. initiated and supervised the study. J.P. engineered SpikeyG and SpikeyGi. J.P. carried out cell culture experiments and analyzed data. X.Y. designed, built, and characterized the ultra-fast two-photon microscope. C.L. developed DeepVID algorithm and analyzed performance. A.M.A. performed animal surgeries for slice and *in vivo* experiments. A.M.A. performed slice experiments and analyzed the data. X.Y. and A.M.A. performed *in vivo* experiments. X.Y., A.M.A., and J.L.C. analyzed *in vivo* data. I.A.C., I.G.D., L.T., V.A.P., and J.L.C. provided input and guidance. J.P., X. Y., A.M.A., C.L., L.T., V.A.P., and J.L.C. prepared the manuscript.

## MATERIALS AND METHODS

### Plasmid construction

The starting constructs used in this study, pcDNA3.1/Puro-CAG-ASAP3b and pcDNA3.1/Puro-CAG-ASAP3b-Kv2.1, were a generous gift from Michael Z. Lin (Villette et al., 2019). The identification of the expressing cells during automated functional screening (see below) was facilitated with the addition of a self-cleaving T2A peptide sequence (GSGEGRGSLLTCGDVEENPGP) followed by nuclear-localized tag-FPs (mCherry) at the C-terminus of GEVI ASAP3. For neuronal expression, GEVI variants SpikeyG and SpikeyGi were subcloned into the pAAV-hSyn-eGFP (Addgene #50465) by replacing eGFP using KpnI and NheI restriction sites. Additionally, a fusion of the C-terminal cytoplasmatic segment of the Kv2.1 channel (the 65AA long proximal restriction and clustering signal) to the C-terminus of the GEVI variants via the GSSGSSGSS linker facilitated restricted expression targeted to the neuronal soma and proximal dendrites (Villette *et al*., 2019). All constructs were manufactured using InFusion Cloning System (Clontech, USA), with all the products confirmed by sequencing (Keck DNA Sequencing Facility, Yale).

### Virus production

AAV2/PhP.eB-hSyn-SpikeyGi-Kv2.1 (9.7×10^12^ gc/mL) was obtained from Boston Children’s Hospital Viral Core. AAV2/8-hSyn-SpikeyGi-Kv2.1 (1.0×10^10^ gc/mL) and AAV2/8-hSyn-SpikeyG-Kv2.1 (1.0×10^10^ gc/mL) were produced and purified in the house using an established protocol (Challis et al., 2019) and commercial purification and titration kits (Takara Bio, USA).

### Library production

The production of GEVI site-directed mutagenic libraries was described previously (Platisa *et al*., 2017, 2020). Briefly, the mutagenic PCR reaction was run with a mix of the three forward primers containing three degenerative codons (NDT, VHG, or TGG) and the single reverse primer. This combination of degenerative primers resulted in site-directed mutagenic libraries encoding all 20 amino acids with repeats for two amino acids, Lysine and Valine. The insertional library that resulted in the development of the SpikeyGi was produced with a single set of PCR primers with the forward primer containing degenerative codon NNK targeted between ASAP3 amino acid residues D151 and N152 (Villette *et al*., 2019). The 15bp long overlapping extension with a sequence identical to a vector at the insertion site in forward primers facilitated vector circularization. For the PCR reaction, we used CloneAmp™ DNA polymerase, and the parent template was removed with *DpnI* restriction enzyme digestion. For the ligation of the amplified mutagenic vectors, we used In-Fusion HD enzyme premix and for bacterial transformation Stellar Chemically competent cells (Clontech, USA). The mutagenic libraries were produced in the 96-well plate with 46 bacterial colonies selected for each (two libraries per 96-well plate; four wells were controls). The cDNA was purified using an automated liquid handling robot (epMotion 5057; Eppendorf, USA), and the library complexity was confirmed by sequencing 10% of selected colonies.

### Cell culture

Functional testing on the semi-automated screening platform was done using spontaneously spiking HEK cells (kind gift from Dr. Adam Cohen of Harvard University; Park *et al*., 2013; #CRL-3269, ATCC, USA) expressing GEVI mutants. The stable expression of NaV 1.3 and KiR 2.1 ion channels creates spike-like electrical activity in these cells. The cells were cultured in DMEM/F12, 10% FBS, 1% penicillin (100 U/mL), streptomycin (100 μg/ml), geneticin (500 μg/mL) and puromycin (2 μg/mL) (Sigma-Aldrich). For simultaneous patch-clamp and imaging GEVI testing, we used HEK 293 cells (# CRL-1573, ATCC, USA). The cells were kept in Dulbecco’s Modified Eagle Medium (DMEM, High glucose; Invitrogen, USA) supplemented with 10% fetal bovine serum (FBS; Sigma-Aldrich, USA). The cultures were maintained in a humidified incubator at 37°C in a 5% CO2 environment, and cells were experimentally used for up to 25 passages. For functional testing, cells were plated either on glass-bottom 96-well black dishes (Cellvis, USA) or on 12-mm coverslips (Carolina Biological, USA) coated with poly-D-lysine hydrobromide (Sigma-Aldrich, USA). For transient expression of GEVI variants, we used Lipofectamine 200 (Invitrogen, USA) at the half of the manufacturer’s recommended reagent amount (for DNA 0.1 ʼg per 96-well or 0.4 μg per 12 mm coverslip in 24-well dish, for Lipofectamine 2000 0.25 ʼl per 96-well or 1 μl per 24-well).

### Functional testing of mutagenic libraries under widefield excitation

The functional screening of GEVI mutants is done as previously described (Platisa et al., 2020; Platisa et al., 2017). In short, a semi-automated screening platform was built around a Nikon Eclipse Ti-E inverted microscope equipped with a Perfect Focus System and a motorized Prior Proscan II stage (Prior Scientific, Inc., USA). The custom-made imaging chamber holds 96-well plates under constant temperature (37°C) and humidity during experiments. For imaging, we used a Nikon Plan Apo 20x 0.75 NA objective (Nikon, Japan), a pE-300 (CoolLED Ltd, U.K.) light source, and an ORCA Flash 4.0 sCMOS camera (Hamamatsu, Japan). For ASAP-based constructs (GFP), we used a 470/40 nm excitation filter, 495 nm dichroic mirror, and 525/25 nm emission filter (# 49002, Chroma Technologies Corp., USA). The nuclear-localized tag protein, mCherry, was visualized with a 560/40 nm excitation filter, 585 nm dichroic mirror, and 630/75 nm emission filter (#49008, Chroma Technologies Corp., USA). For field stimulation (Grass S48 Stimulator, USA), we used a custom-made field electrode and actuator system (Thorlabs, USA) attached to the roof of the imaging chamber. The image collection, electrical stimulation, and signal detection were done using a custom application written in LabView (National Instruments, Inc., USA). The fluorescence intensity of nuclear-localized mCherry was used to identify expressing cells and select for the field of views within each well. For functional screening, images were collected at 100 fps in 2500 ms long sweeps with a single pulse of 70V 0.5 ms applied at 400 ms from the beginning. The fluorescence signal in each cell in response to field stimulation was quantified as ΔF/F. Each GEVI variant was screened in four separate plates, and the selection of the best mutants was based on the maximum response amplitude across cells and wells.

### Electrophysiology and widefield imaging in HEK293 cells

For whole-cell patch-clamp experiments, HEK239 cells were kept in a perfused chamber at 33-35°C (Warner Instruments, USA) with the constant running bath solution (129 mM NaCl, 4 mM KCl, 1 mM CaCl2, 1 mM MgCl2, 10 mM D-glucose, and 10 mM HEPES, pH 7.4 and was adjusted to 310 mOsm with sucrose). The 3-5 MΩ glass patch pipettes (capillary tubing with 1.5/0.75 mm O.D./ID, WPIP, USA) were pulled on a P-97 Flaming/ Brown type micropipette puller (Sutter Instrument Company, USA). The pipette solution contained 125 mM K-gluconate, 8 mM NaCl, 0.6 mM MgCl2, 0.1mM CaCl2, 1 mM EGTA, 4 mM Mg-ATP, 0.4Na-GTP and 10 mM HEPES, pH 7.2 and adjusted to ∼290 mOsm. Voltage-clamp recordings in the whole-cell configuration were performed using MultiClamp 700B amplifier, digitizer Digidata Series 1400A and pClamp software (Molecular Devices, USA). Depending on the experiment, the voltage was changed from a holding potential of -70mV to i) +30 mV (a single-step depolarization experiments), ii) -110, -90, -50, -30. -10, 10, and 30mV) (in a series of 20mV incremental subsequent steps), or iii) -120, -20, +30 and +80 mV (a series of steps). Imaging was performed on an Olympus BX61WI upright microscope using a LUMPlan FL 40x N.A. 0.80 water immersion objective (Olympus, USA) and a 488 nm 50 mW laser light excitation (DL488-050, USA). We used a GFP filter set, a 495 nm dichroic mirror, and a 520/35 nm emission filter (Semrock, USA), and the laser power measured at the preparation was 13-18 mW/mm2. Additionally, the light intensity was adjusted for each recording session using a continuous circular neutral density filter (ThorLabs, Inc., USA) to the minimum required to record optical signals. The images were collected with a fast-speed NeuroCCD camera controlled by NeuroPlex software (RedShirt Imaging, USA) at a frame rate of 500 or 1000Hz. For the image demagnification, we used either an Optem zoom system A45731 0.13 or Optem C-to-C mount 25-70-54 0.383 (Qioptiq LINOS, USA).

### Widefield imaging analysis

The data were analyzed using NeuroPlex, Excel, and custom scripts written in Igor and Matlab. All the results are presented as a mean value and the standard error of the mean (s.e.m.). The values for the resting fluorescence and the bleaching rate were derived from recordings on the screening platform. The resting fluorescence was calculated as the mean value recorded across all the cells within the field of view at the first five frames at the beginning of the recording and prior stimulation. The bleaching rate is calculated as a percent change between resting fluorescence at the beginning and end of the trial. The optical traces are spatial averages of the intensity of the pixels within the region of interest (ROI) that covers the cell body. The ROIs were visually identified using the Neuroplex feature Frame Subtraction. The amplitude of fluorescence change was measured as the difference between the averaged values for 50 frames before stimulation and 5 (for SpikeyGi) and 100 (ASAP3 and SpikeyG) frames around the peak of the response. Data are presented as the voltage-dependent change in fluorescence divided by the resting fluorescence, ΔF/F. For bleach correction, the portion of the trace outside the stimulus was fitted with a double exponential curve.

### Animal preparation

All experimental procedures were approved by the Institutional Animal Care and Use Committee for the Charles River Campus at Boston University. For *in vitro* slice experiments, viral injections were performed in neonatal (P7-P9) C57Bl/6 mice. Mice were bred in house in the Boston University animal care facility, with standard housing, a 12-hour light/dark cycle, and *ad lib* access to food and water. Pups were removed from the mother, anesthetized with 1-3% isoflurane, and placed in a custom stereotaxic holder for neonatal mice. A small incision was made in the scalp and a two small holes were made with a 30-gauge needle in the skull of the left hemisphere. Mice were injected with AAV2/8-hSyn-SpikeyGi-Kv2.1 or AAV2/8-hSyn-SpikeyG-Kv2.1 (300 nL) at 250 μm below the pial surface. Incisions were closed with Vetbond Tissue Adhesive (3M), and animals were treated post-operatively with buprenorphine (0.05-1 mg/kg, s.c.). Animals were returned to their mother immediately after surgery and weaned at 3 weeks of age.

For *in vivo* experiments, adult (6-8 week old) C57Bl/6 mice, stereotaxic viral injections of GEVIs were performed in L2/3 and L5 of primary somatorsensory cortex, 300 and 500 μm below the pial surface (600 nL total volume). Either AAV2/PhP.eB-hSyn-SpikeyG-Kv2.1 (1:8 or 1:12 in saline) or AAV2/8-hSyn-SpikeyG-Kv2.1 were injected. There were two injection sites per mouse, both targeting the left hemisphere sensorimotor (S1) cortex (AP -1.1 mm, ML 2.9 mm and AP -1.1 mm, ML 3.3 mm). To enable optical access, a 4 mm cranial window was implanted over S1 (Margolis et al., 2012). A metal headpost was implanted over the right hemisphere to enable head fixation. Animals were treated post-operatively with buprenorphine (0.05-1 mg/kg, s.c.) and given 2-3 weeks for viral expression to take place. Animal were handled and habituated to head fixation for 3-5 days before imaging experiments began.

### Slice electrophysiology

For *in vitro* characterization of GEVIs, whole-cell slice electrophysiology was performed on layer 2/3 cortical neurons. To obtain brain tissue slices, mice were deeply anesthetized with ketamine/xylazine and trancardially perfused with modified ACSF (in mM, 124 NaCl, 75 sucrose, 10 glucose, 2.5 KCl, 1.25 NaH_2_PO_4_, 25 NaHCO3, 1.3 ascorbic acid, 0.5 CaCl_2_ and 7 MgCl_2_). Brain tissue was sliced into 300 μm coronal sections and maintained with oxygenated (95/5% O2/CO2) ACSF (in mM, 124 NaCl, 26 NaHCO3, 20 sucrose, 3 KCl, 1.25 NaH2PO4, 2 CaCl2 and 1.5 MgCl_2_). Pipettes were pulled from thin-walled borosilicate capillaries without filament (3-6 MΩ) (Sutter Instruments), and filled with internal solution containing (in mM, 135 K-gluconate, 10 HEPES, 2 MgCl_2_, 2 MgATP, 0.4 EGTA, 0.5 Na_3_GTP, 10 phosphocreatine disodium). The process of approaching the cell, sealing onto the membrane, and breaking in to establish a whole-cell configuration was controlled by Autopatcher (Neuromatic Devices). Electrical signals were recorded with a Multiclamp 700B amplifier (Axon Instruments), filtered at 10 kHz and digitized at 20 kHz (National Instruments PCI-6321), and recorded with custom MATLAB software (Mathworks). Simultaneous 2P imaging was performed at with a conventional 2P microscope (Ultima, Prairie Technologies, Middleton WI) using a 40x/0.8NA water immersion objective (Olympus) and Dodt contrast imaging. For laser source, a 80Mhz Ti:sapphire laser (Mai Tai HP, Spectra Physics) was tuned to 920nm with 35-45mW delivered at the sample.

For characterization of fluorescence responses to varying membrane potentials, fluorescence intensity was measured while neurons were held in voltage clamp mode. One-second trials consisted of a 200ms baseline period at resting membrane potential (−70 mV), followed by a 400 ms voltage step, then a return to resting membrane potential for 400ms. Trials were repeated 10 times across 10 voltage steps ranging from -130 mV to +50 mV (20 mV increments). For image acquisition, frame scans of the targeted cell were acquired at 20Hz across 128 × 85 pixels at ∼0.3μm/pixel resolution and 2.8ʼs pixel dwell time. For analysis, the cell membrane was segmented by isolating pixels 1.3-2.0x above background. For characterization of action potential responses, neurons were recorded in current clamp mode with single action potentials evoked at 100ms intervals (triggered by 10ms pulses of 800-1200 pA), with 10 500-ms sweeps of 5 action potentials each. Fluorescent responses were recorded by line scans along the cell membrane at 24-54 pixels per line, scanned at 3-5 kHz, 2.8ʼs pixel dwell time. ΔF/F_0_ was calculated where F_0_ corresponded to fluorescence at resting membrane potential.

### Microscope design

The optical design was performed using Zemax OpticStudio software (Zemax LLC), and the opto-mechanical design was performed using AutoDesk Inventor (Autodesk Inc.). A 920nm, 2W fiber laser (ALCOR-920, Sparks Lasers, 100fs pulse duration, 31.25MHz repetition rate) was used as the light source. The total laser power was controlled by a Pockel’s Cell (350-80, ConOptics). The laser beam was split into 4 beam paths using polarizing beamsplitter (PBS) cubes (PBS123/ CCM1-PBS253, Thorlabs). A half-wave plate (AHWP05M-980, Thorlabs) was placed before each beamsplitter cube (three in total) to allow power control of individual paths. Two half-wave plates were mounted on manual rotation mounts (CRM1PT/M, PRM05/M, Thorlabs) The third half-wave plates mounted on a motorized precision rotation stage (PRM1/MZ8, Thorlabs) for remote control when selecting between 8 beam and 1 beam imaging. Temporally multiplexed beams were delayed by 8ns relative to each other using delay lines. After the delay line, all beams were reduced to obtain a 1.5mm beam diameter (BE052-B, AC127-075-B, ACN127-025-B, or BE02-05-B, Thorlabs).

Temporally multiplexed beams were routed into customized beamsplitter plates. Beamsplitter plates serve two functions. First, they split each temporally multiplexed beam into pairs of spatially multiplexed beams. Second, they arrange the beamlets linearly to definite the position of each subarea. A beamsplitter plate consisted of an assembly of half-wave plates (AHWP05M-980, Thorlabs) on rotation mounts (PRM05/M, Thorlabs), PBS cubes (PBS123, PBS053, Thorlabs) and half-inch gold-coated mirrors (PF05-03-M01, Thorlabs) on miniature mirror holders (LMMH-12.7R-N, OptoSigma). One beamsplitter routed 4 beams through independent f=100mm relay lenses (KPX034AR.16, Newport) which were held in a customized lens holder and separated by 8.5mm, corresponding to 100μm spacing at the sample. Two beamsplitter plates were used to combine a total of 8 beams using a 2-inch PBS cube (PBS513, Thorlabs) and positioned such that beams from each beamsplitter plate were interleaved at 50μm spacing at the sample. A beam blocker mounted onto a linear actuator (L12-P, Actuonix) was placed after each lens holder and allowed selection between all 8 beams or any arbitrary individual beam.

All 8 beamlets were sent to a customized f=330mm scan lens assembly (SLB-50-300N, SLB-50-450P, SLB-50-250P, OptoSigma), and conjugated to a 2kHz resonant scanner (CRS 12kHz, 5.0mm x 7.2mm aperture, Cambridge Technology) and galvo scanner (6215H, Cambridge Technology). The scan lens (S4LFT0089/094, Sill Optics GmbH & Co.) and tube lens (AC508-500-B, AC508-750-B, Thorlabs) expanded the beam to fill the back aperture of the objective (N16XLWD-PF, 16x, Nikon). The laser light was reflected to the objective by a short-pass filter (FF01-720/SP-25, AVR Optics).

The emitted light passed through the short-pass filter and then was separated by a secondary dichroic (FF556-SDi01-40×45, Semrock) into green and red fluorescent channels. Red fluorescence excited from a single beam was detected using an achromatic lens (AC508-080-A, Thorlabs) and an eyepiece (19mm, Televue) focused onto a hybrid PMT (R11322U-40, Hamamatsu). For green fluorescence, a customized lens assembly was used to magnify and reshape the square FOV to match the geometry of a 16×1 multi-anode photomultiplier (H13123, GaAsP 16-channel MAPMT, Hamamatsu). The assembly included two achromatic lenses (AC508-100-A, Thorlabs) and two cylindrical lenses (LJ1567RM-A, Thorlabs, and CKX019, Newport) positioned orthogonal to each other.

Using a customized 16-to-4 adder (Marina Photonics Inc.), signals from the 16 MAPMT anodes were amplified, digitized, and then combined into four detector subgroups each consisting of signals from 4 adjacent anodes. Using custom electronics, the subsequent LVPECL signal was converted and fanned out to two LVDS signals (NB6N11S, ON Semiconductor). The signals were collected by an FPGA (PXIe-7965R, National Instruments Corp.) with a 20-channel digital I/O board (NI-6587, National Instruments Corp.) operating at 1Gbit/s sampling rate. For a given detector subgroup, the two fanned out signals were inputted into two I/O channel wherein one of the two fanned out signals was delayed by 0.5ns. This provided a 2Gbit/s sampling rate for each detector subgroup when combined on the FPGA.

Each detector subgroup contained fluorescence from two beamlets which were then temporally demultiplexed on the FPGA using a synchronization signal provided by the laser and electronic delay box (DB64, Stanford Research Systems) providing de-multiplexing with 0.5ns precision. Additional time-dependent gating was implemented on the FPGA to minimize spatial multiplexed crosstalk from neighboring detector subgroups (LabVIEW, National Instruments). For sample positioning, a lifting stage (HT160-16-DC, Steinmeyer Mechatronik GmbH) mounted on an XY stage (KT310-200-DC, Steinmeyer Mechatronik GmbH) was used. The microscope system was controlled by a customized C++-based software, Scope. The software controlled the resonant and galvo scanner (imaging FOV), the pockel’s cell (laser power) and the shutter through a DAQ module (PXI-6259, National Instruments Corp.).

### Point spread function characterization

For each beamlet in the UF2P microscope, the point spread function (PSF) was measured using 0.5μm diameter fluorescent beads with (T7281, Invitrogen). Images were taken one beamlet at a time. The beads were embedded in 1.5% agarose and an image stack of the bead was taken at 0.1×0.1×0.5μm^3^ voxel resolution with 50 frames at each plane. Motion correction was performed at each plane before averaging. A summed Z-intensity projection of the image stack was taken and the maximum intensity pixel was identified as the center of the bead along the X/Y-axis. The center along the Z-axis was defined as the plane with the maximum total signal. The profile was plotted through the center along each axis. The PSF value reported was the full-width-at-half-maximum (FWHM) of each profile.

### Crosstalk characterization

To determine the degree of temporal and spatial crosstalk observed in scattering tissue, a mouse virally expressing SpikeyGi implanted with a cranial window was used. Tissue was scanned using one beam at a time (∼30mW) and fluorescence was collected across all 8 subareas. Images consisting of 50-frame averages acquired at 389Hz were taken at 30μm steps from 0-300μm below the pial surface. For the excited subarea in each image, pixels corresponding to SpikeyGi fluorescence were identified as those whose intensities were >95^th^ percentile compared to other pixels in the same subarea. Percent crosstalk was calculated as the mean signal measured in each of the non-excited subareas divided by the mean signal in the excited subarea contain SpikeyGi fluorescence.

### Intrinsic signal optical imaging

To identify the location of viral expression relative to S1, intrinsic signal imaging was performed under light anesthesia (1-1.5% isoflurane). The cortical surface was illuminated with a 625 nm LED (Thor Labs), and two individual whiskers (B2 or C2) were stimulated at 10 Hz with a piezo-electric stimulator. Reflectance images were recorded using a f = 25 mm lens (Navitar) and a CMOS camera with a 30 Hz frame rate, 6.5 μm pixel size, 4 × 4 binning, and 512 × 512 binned pixels (Hamamatsu). Cortical activation in the barrel column was determined by comparing changes in reflectance during whisker stimulation versus periods of non-stimulation, expressed as ΔR/R_0_ (150 frame average). Barrel columns were identified as signal minima after averaging intrinsic reflectance signals over 10 trials.

### Self-supervised deep learning denoising of voltage imaging data

DeepVID combines self-supervised frameworks implemented in DeepInterpolation and Noise2Void (Lecoq et al., 2021; Li et al., 2021). The network architecture of DeepVID was based on the DnCNN (Zhang et al., 2017), a fully convolutional network with residual blocks. This architecture was chosen to better accommodate the 8:1 aspect ratio in the sub-image scanned by each beamlet. The network was constructed with 2D convolution layers (Conv), batch normalization (BN) layers and Parametric Rectified Linear Unit (PReLU) activation layers, with 16 repeated residual blocks in the middle. Each residual block contained two 3×3 Conv layers with BN layers followed, and an PReLU activation layer was appended after the first BN layer. The skip connection was added to link low dimensional and high dimensional features by adding the feature map of the input and the output for each residual block (**Figure 4A**).

DeepVID was designed to denoise a single frame from each sub-area at a time. It was trained to predict the central frame *N*_*0*_ using an input image time series, consisting of *N*_pre_ frames before and *N*_post_ frames after the central frame, in addition to a degraded central frame with several “blind” pixels. A random set of pixels (*p*_blind_) in the central frame were set as blind pixels using a binary mask, whose intensities were replaced by a random value sampled from randomly selected pixels in the frame. The hyperparameters (*N*_pre_= 3, *N*_post_ = 3, p_blind_ = 10%) were optimized to maintain the temporal dynamic of voltage signal spikes while recovering a high single-frame spatial resolution. The loss function was the mean squared error (i.e. L2 loss) between the original and denoised central frame and was calculated only on the blind pixels. The training was performed using the Adam optimizer with 360 steps per epoch and a batch size of 4. The training stopped after going over all samples in the data set one time to avoid overfitting. The learning rate was initialized at 5×10^−6^ and reduced to 1×10^−6^ when the loss on the validation set did not decrease in the past 288,000 samples.

The training data set consisted of 1181 videos, each of which contained 1000 frames acquired at 803 Hz. Each image time series was preprocessed by detrending and normalization before sending into the DeepVID network. For detrending, the trend was approximated by a second order estimator by fitting to the time trace of the intensity mean of each frame. The scalar in the trend at each frame was subtracted from all the pixels in the corresponded frame. The detrended video was then normalized by subtracting the mean and dividing by the standard deviation of all pixels in the image time series. The network was trained on a graphics processing unit (Nvidia P100, 12 GB VRAM) using TensorFlow 2.2.0, Python 3.8 and CUDA

11.2. Once DeepVID was trained, inference denoising of subsequent image data using the trained model was performed frame-by-frame by feeding each corresponding 7-frame image time series. It can be performed at approximately 200 frames per second on a single Nvidia P100 GPU. To quantify the performance of DeepVID, the single-pixel SNR was defined as the ratio of the mean divided by the standard deviation of the pixel intensity along the temporal axis.

### *In vivo* voltage imaging

To measure sensory-evoked voltage responses, whisker stimulation was performed on awake, head-fixed mice. Air puffs (25ms) were delivered to the whisker pad contralateral to the imaged hemisphere. Stimulus patterns consisted of alternating trains of 1 puff or 5 puffs separated at 4 second intervals. Trains of 5 air puffs were delivered at either 5 or 10 Hz stimulus frequency. Using the UF2P microscope, voltage imaging in S1 was performed 803Hz at frame rate across 400×192 total pixels (400×24 per subarea) at x: 1.0 μm/pixel, y: 2.1 μm/pixel resolution, ∼30mW per beamlet, ∼0.1ʼs dwell time. For image analysis, each subarea was analyzed independently in MATLAB (Mathworks). For correction of brain motion, brain motion was estimated using a 2D rigid fast-Fourier transform based on images from one subarea and then a rigid transform correction was applied to all 8 subareas. Images for each subareas were then denoised using DeepVID. Neurons were then manually segmented and fluorescence trace extracted for each ROI.

### Action potential analysis

For analysis and detection of spiking-related voltage signals, slow fluctuations in fluorescence signals for each cell were first removed by baseline subtraction along a moving average across 2.5 seconds of recordings. Given the ∼5ms rise and ∼12.5ms decay time (peak-to-trough) of tested GEVIs, a putative spike trace was generated by calculating the difference in fluorescence intensity across every 10^th^ imaging frame (12.5ms) across the time series. The putative spike trace was normalized by the cell’s noise level, defined as the mean of the absolute difference between each time point, to produce an SNR trace. Spikes were identified as transient events exceeding a given SNR threshold. Sensory-evoked action potentials were identified as detected spikes occurring within a 20 frame window (25ms) following air puff delivery. The sensitivity of GEVIs for spike detection was assessed by quantifying the percent of detected action potentials evoked for each air puff across a range of SNR thresholds.

### Photobleaching

For *in vitro* photobleaching measurements, cortical tissue was prepared in the same manner and imaged using the same setup for slice electrophysiology. Areas of SpikeyGi or SpikeyG expression were positioned in the field of view using an epifluorescent microscope. Imaging data was recorded at 1 Hz in an intermittent pattern with 10s on (shutter open) and 5s off (shutter closed) for 12.5 min. Laser power of 35 mW was used, with 1.13 μm per pixel and dwell time of 2.8 ʼs. ROIs corresponding to a single cell were extracted manually, and each data point corresponded to the mean fluorescence intensity in a single frame.

*In vivo* photobleaching measurements were performed using the UF2P microscope on awake, head-fixed mice with implanted cranial windows. Imaging was performed using a total laser power of 240mW at the sample (30mW per beam). The tissue was scanned at a framerate of 803Hz in an intermittent pattern consisting of 9s on (shutter open) and 4s off (shutter closed) for 60 min. To assess action potential responses during photobleaching, voltage imaging with whisker stimulation was performed at 15 minute intervals. ROIs corresponding to a single cell were extracted manually. Each data point corresponded to the mean fluorescence intensity of every 20^th^ frame across the 9 seconds of imaging. Photobleaching rates were determined by normalizing all data points to the first data point.

### Photodamage

Animals previously injected with virus and implanted with cranial windows were used for experiments. A location for laser exposure away from the viral injection sites was selected based on a wide-field blood vessel map under the cranial window. Each photo-damage session consisted of one-hour of continuous laser scanning with 240mW total power (30mW per beamlet). Sixteen hours after laser exposure, animals were anesthetized with 1.5-3.0% isoflurane and cranial windows were removed to expose the cortex. The blood vessel map was used to locate the laser exposure site, and a lipophilic dye (SP-DiIC_18_(3); ThermoFisher Scientific; D7777) was injected to mark the four corners of the laser exposure field of view. Animals were then transcardially perfused with 0.1 M PBS and 4% paraformaldehyde. Brains were postfixed in 4% paraformaldehyde for 24 hours, transferred to 0.1 M PBS, then were sliced with a vibratome into 50-μm coronal sections. Slices were first incubated in a blocking solution (10% normal goat serum and 1% Triton X-100) and washed three times in 0.1M PBS. Alternating slices were labeled with sets of primary antibodies in 5% normal goat serum and 0.1% Triton X-100. One set of slices were stained with primary antibodies for mouse monoclonal anti-GFAP (G3893; Sigma-Aldrich; 1:1,000 dilution) and rabbit anti-Iba1 (019-19741; Wako Chemicals; 1:500 dilution). The other set of slices were stained with mouse anti-HSP70/HSP72 (ADI-SPA-810-D; Enzo Life Sciences; 1:400 dilution), and rabbit anti-cleaved caspase-3 (Asp175) (9661; Cell Signaling; 1:250 dilution). Slices were then washed three times in 0.1 M PBS and incubated in secondary antibodies. Iba1 and caspase-3 were labelled with goat anti-rabbit Alexa Fluor 555 (Invitrogen, A21429; 1:500 dilution), and GFAP and HSP were labelled with goat anti-mouse Alexa Fluor 647 (Invitrogen, A21235; 1:500 dilution). Slices were washed in 0.1M PBS and mounted with Fluoromount-G mounting medium (0100-01, SouthernBiotech). The lipophilic dye was used to identify which sections contained the laser exposure site, and where it was located on the medial-lateral axis. Slices were imaged with a Nikon ECLIPSE Ni-E microscope and NIS-Elements software (Nikon Instruments).

Photodamage was determined increased antibody labeling in the laser exposed region relative to the corresponding area on the contralateral hemisphere. The relative fluorescence was determined by dividing the mean fluorescent intensity (a.u.) on the treated hemisphere by mean intensity on the contralateral hemisphere. Fluorescence was measured in two areas of the cortex, the laser exposure site and a control area that was at least 1 mm away from the site of laser exposure. The control area was intended to capture protein expression that may have been caused by chronic window implantation and/or virus injection, but was not caused by laser exposure.

### Material and data availability

All sequence information will be available at NCBI, all plasmids and rAAVs will be available through Addgene and UNC Vector Core. All codes will be available at https://github.com/common-chenlab/ and https://github.com/bu-cisl/DeepVID

## SUPPEMENTARY INFORMATION

**Figure S1 related to Figure 3.**
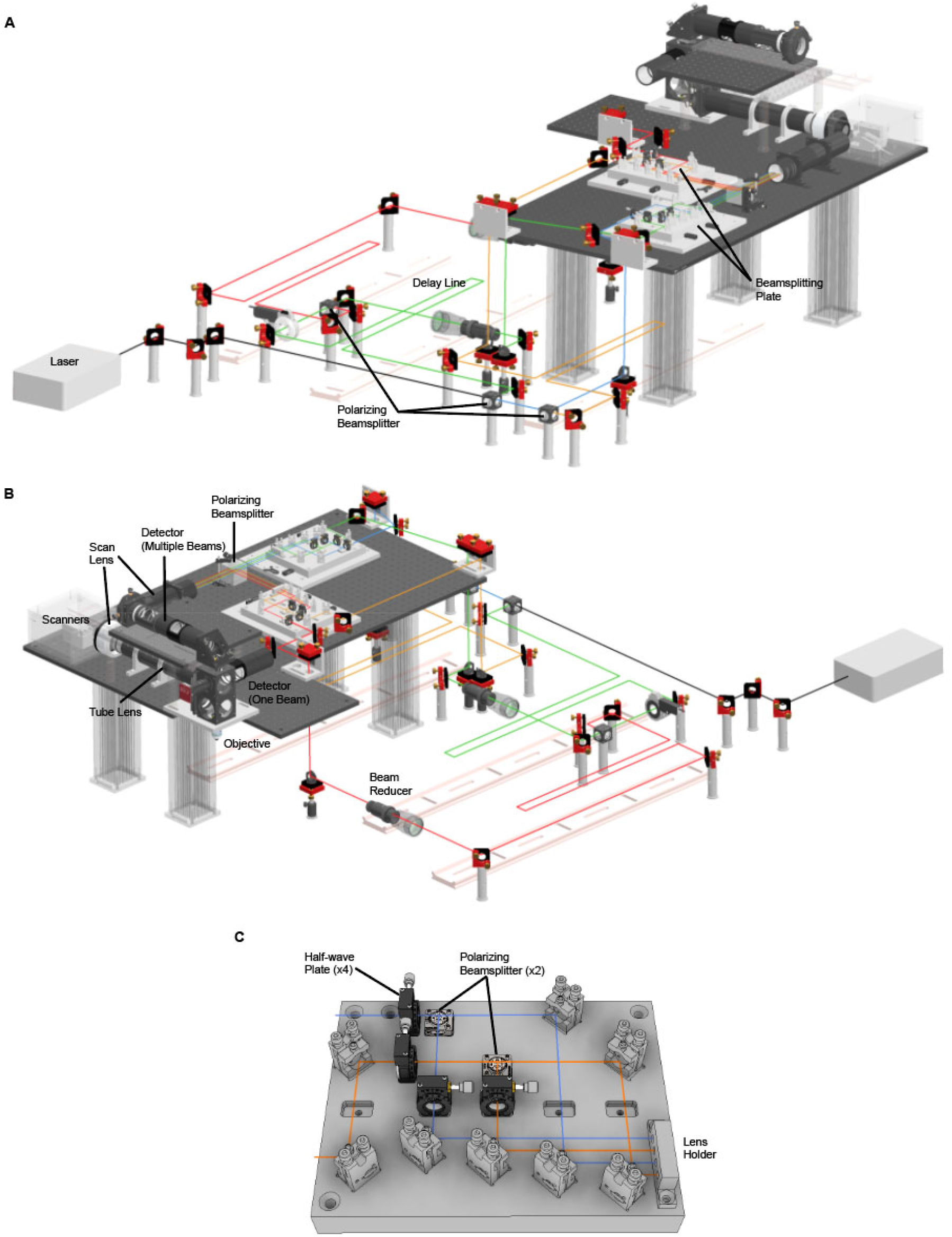
Opto-mechanical design of UF2P microscope. **(A-B)** CAD design Rendering of the Uf2P microscope. **(C)** CAD design rendering of the beam splitter plate.

**Figure S2 related to Figure 3.**
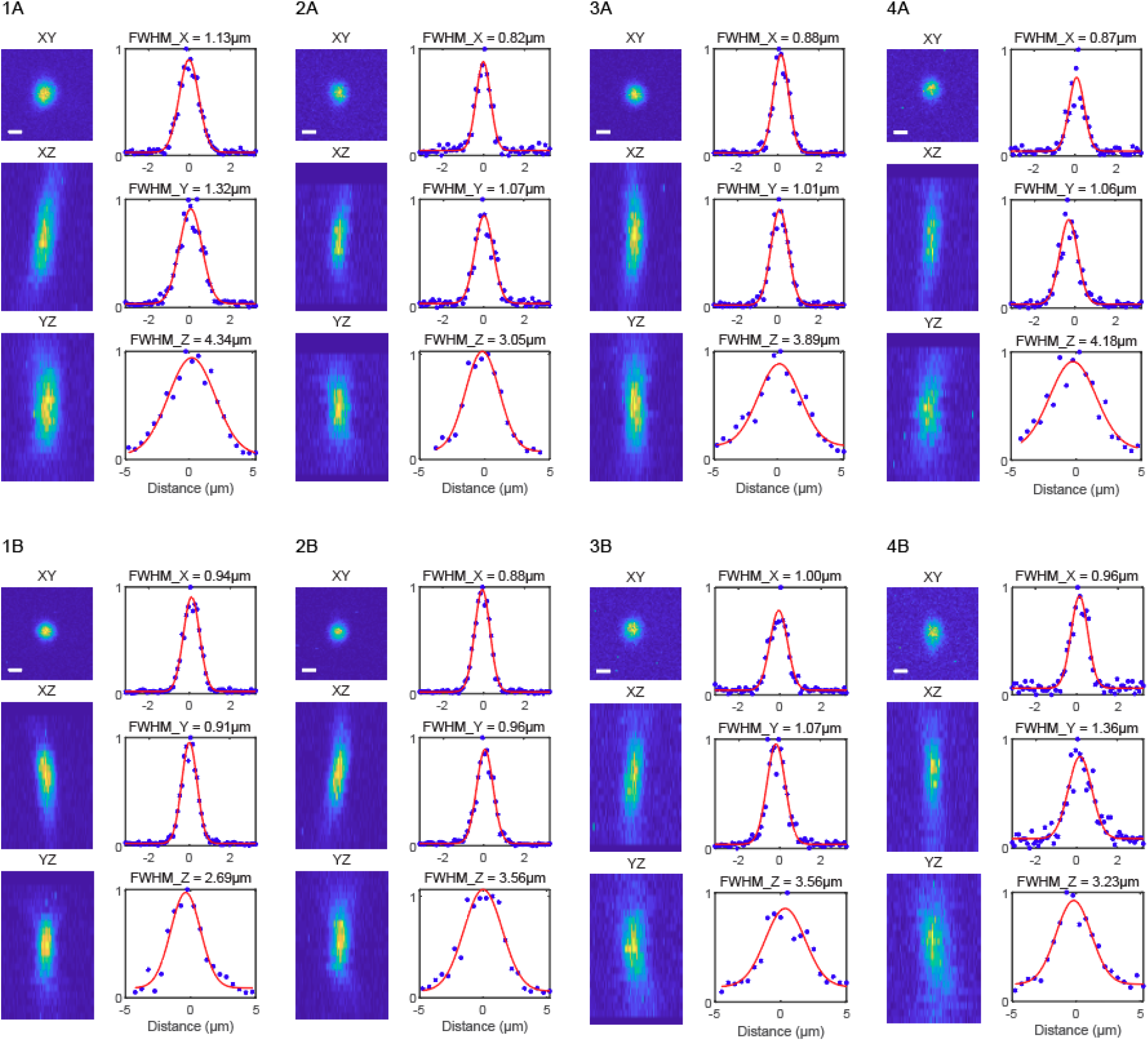
Example point spread function measurements for each beamlet in the UF2P microscope.

**Figure S3 related to Figure 5.**
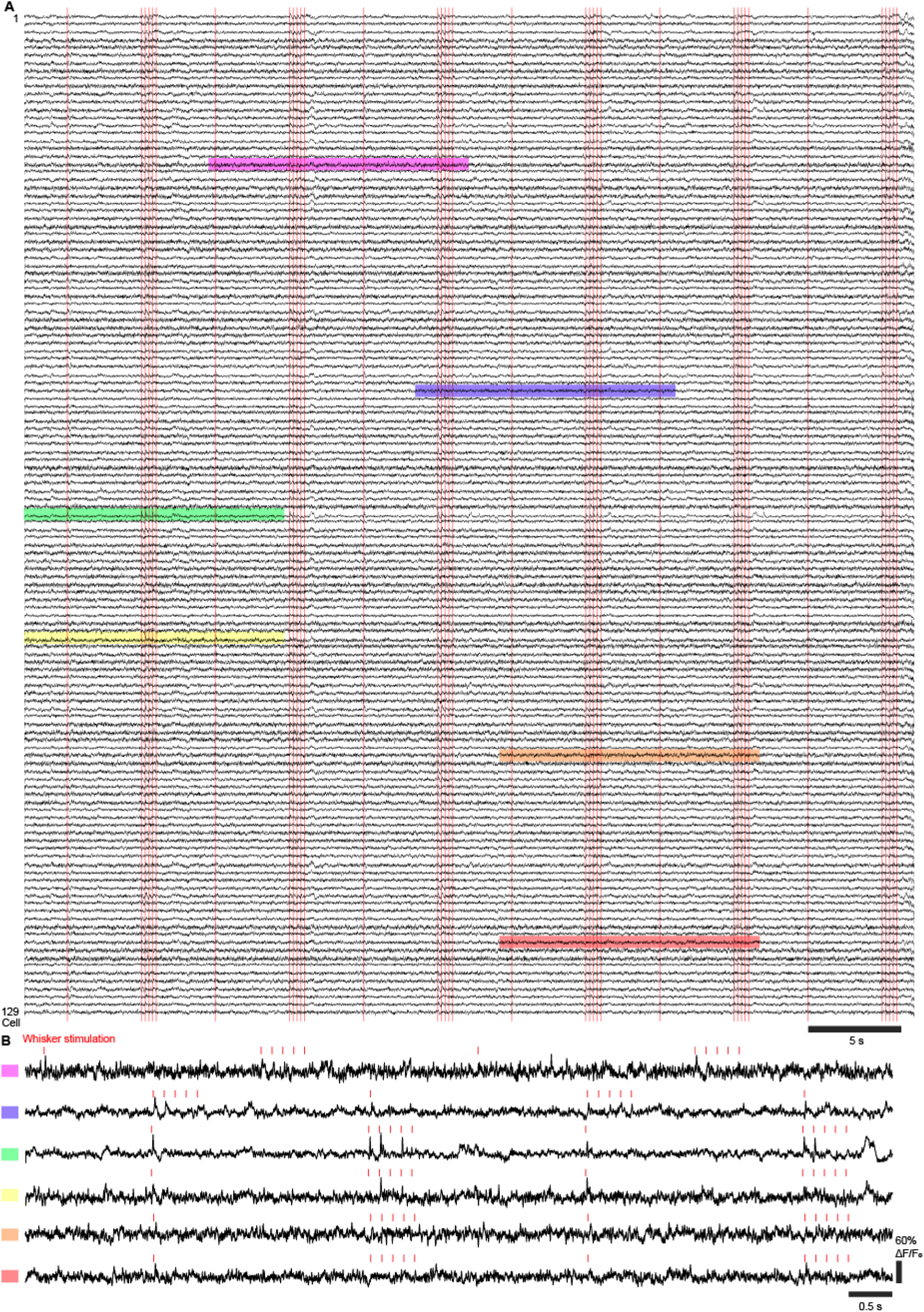
*In vivo* population imaging of SpikeyGi from UF2P microscope. **(A)** Fluorescence traces from simultaneous recordings across 129 S1 neurons from the UF2P microscope acquired at 803 Hz. Raw images were denoised with DeepVID. For visualization purposes, traces are detrended with 2.5 sec moving average and low pass filtered at 200 Hz. Red lines indicate air puff whisker stimulus. Detected action potentials are plotted in Figure 5B. **(B)** Magnified view of example traces across different cells. Squares denote traces corresponding to shaded regions of the same color indicated in [A]. Red lines indicate air puff whisker stimulus.

## Notes

### Competing Interest Statement

The authors have declared no competing interest.

## REFERENCES

Abdelfattah, A.S., Kawashima, T., Singh, A., Novak, O., Liu, H., Shuai, Y., Huang, Y.-C., Campagnola, L., Seeman, S.C., Yu, J., et al. (2019). Bright and photostable chemigenetic indicators for extended in vivo voltage imaging. Science 365, 699–704.

Abdelfattah, A.S., Valenti, R., Zheng, J., Wong, A., Team, G.P., Podgorski, K., Koyama, M., Kim, D.S., and Schreiter, E.R. (2020). A general approach to engineer positive-going eFRET voltage indicators. Nat Commun 11, 3444.

Adam, Y., Kim, J.J., Lou, S., Zhao, Y., Xie, M.E., Brinks, D., Wu, H., Mostajo-Radji, M.A., Kheifets, S., Parot, V., et al. (2019). Voltage imaging and optogenetics reveal behaviour-dependent changes in hippocampal dynamics. Nature 569, 413–417.

Amir, W., Carriles, R., Hoover, E.E., Planchon, T.A., Durfee, C.G., and Squier, J.A. (2007). Simultaneous imaging of multiple focal planes using a two-photon scanning microscope. Opt Lett 32, 1731–1733.

Bando, Y., Sakamoto, M., Kim, S., Ayzenshtat, I., and Yuste, R. (2019). Comparative Evaluation of Genetically Encoded Voltage Indicators. Cell Rep 26, 802–813 e804.

Chamberland, S., Yang, H.H., Pan, M.M., Evans, S.W., Guan, S., Chavarha, M., Yang, Y., Salesse, C., Wu, H., Wu, J.C., et al. (2017). Fast two-photon imaging of subcellular voltage dynamics in neuronal tissue with genetically encoded indicators. Elife 6.

Charan, K., Li, B., Wang, M., Lin, C.P., and Xu, C. (2018). Fiber-based tunable repetition rate source for deep tissue two-photon fluorescence microscopy. Biomed Opt Express 9, 2304–2311.

Chen, J.L., Voigt, F.F., Javadzadeh, M., Krueppel, R., and Helmchen, F. (2016). Long-Range population dynamics of anatomically defined neocortical networks. Elife 5, e14679.

Cheng, A., Goncalves, J.T., Golshani, P., Arisaka, K., and Portera-Cailliau, C. (2011). Simultaneous two-photon calcium imaging at different depths with spatiotemporal multiplexing. Nature methods 8, 139–142.

Clough, M., Chen, I.A., Park, S.W., Ahrens, A.M., Stirman, J.N., Smith, S.L., and Chen, J.L. (2021). Flexible simultaneous mesoscale two-photon imaging of neural activity at high speeds. Nat Commun 12, 6638.

Demas, J., Manley, J., Tejera, F., Barber, K., Kim, H., Traub, F.M., Chen, B., and Vaziri, A. (2021). High-speed, cortex-wide volumetric recording of neuroactivity at cellular resolution using light beads microscopy. Nat Methods 18, 1103–1111.

Feldmeyer, D., Brecht, M., Helmchen, F., Petersen, C.C., Poulet, J.F., Staiger, J.F., Luhmann, H.J., and Schwarz, C. (2012). Barrel cortex function. Progress in neurobiology.

Huang, L., Ledochowitsch, P., Knoblich, U., Lecoq, J., Murphy, G.J., Reid, R.C., de Vries, S.E., Koch, C., Zeng, H., Buice, M.A., et al. (2021). Relationship between simultaneously recorded spiking activity and fluorescence signal in GCaMP6 transgenic mice. Elife 10.

Jin, L., Han, Z., Platisa, J., Wooltorton, J.R., Cohen, L.B., and Pieribone, V.A. (2012). Single action potentials and subthreshold electrical events imaged in neurons with a fluorescent protein voltage probe. Neuron 75, 779–785.

Kazemipour, A., Novak, O., Flickinger, D., Marvin, J.S., Abdelfattah, A.S., King, J., Borden, P.M., Kim, J.J., Al-Abdullatif, S.H., Deal, P.E., et al. (2019). Kilohertz frame-rate two-photon tomography. Nat Methods 16, 778–786.

Kim, K.H., Buehler, C., Bahlmann, K., Ragan, T., Lee, W.-C.A., Nedivi, E., Heffer, E.L., Fantini, S., and So, P.T.C. (2007). Multifocal multiphoton microscopy based on multianode photomultiplier tubes. Opt Express 15, 11658–11678.

Krull, A., Buchholz, T.-O., and Jug, F. (2018). Noise2Void - Learning Denoising from Single Noisy Images. 181110980 [cs].

Lecoq, J., Oliver, M., Siegle, J.H., Orlova, N., Ledochowitsch, P., and Koch, C. (2021). Removing independent noise in systems neuroscience data using DeepInterpolation. Nat Methods 18, 1401–1408.

Li, X., Zhang, G., Wu, J., Zhang, Y., Zhao, Z., Lin, X., Qiao, H., Xie, H., Wang, H., Fang, L., et al. (2021). Reinforcing neuron extraction and spike inference in calcium imaging using deep self-supervised denoising. Nat Methods 18, 1395-1400.

Piatkevich, K.D., Bensussen, S., Tseng, H.A., Shroff, S.N., Lopez-Huerta, V.G., Park, D., Jung, E.E., Shemesh, O.A., Straub, C., Gritton, H.J., et al. (2019). Population imaging of neural activity in awake behaving mice. Nature 574, 413–417.

Platisa, J., Han, Z., and Pieribone, V.A. (2020). Different categories of fluorescent proteins result in GEVIs with similar characteristics. bioRxiv, 2020.2005.2006.081018.

Platisa, J., Vasan, G., Yang, A., and Pieribone, V.A. (2017). Directed Evolution of Key Residues in Fluorescent Protein Inverses the Polarity of Voltage Sensitivity in the Genetically Encoded Indicator ArcLight. ACS Chem Neurosci 8, 513–523.

Podgorski, K., and Ranganathan, G. (2016). Brain heating induced by near-infrared lasers during multiphoton microscopy. J Neurophysiol 116, 1012–1023.

Sjulson, L., and Miesenbock, G. (2007). Optical recording of action potentials and other discrete physiological events: a perspective from signal detection theory. Physiology (Bethesda) 22, 47–55.

Vaswani, A., Shazeer, N., Parmar, N., Uszkoreit, J., Jones, L., Gomez, A.N., Kaiser, Ł., and Polosukhin, I. (2017). Attention is all you need. Paper presented at: Advances in neural information processing systems.

Villette, V., Chavarha, M., Dimov, I.K., Bradley, J., Pradhan, L., Mathieu, B., Evans, S.W., Chamberland, S., Shi, D., Yang, R., et al. (2019). Ultrafast Two-Photon Imaging of a High-Gain Voltage Indicator in Awake Behaving Mice. Cell 179, 1590-1608.e1523.

Weisenburger, S., Tejera, F., Demas, J., Chen, B., Manley, J., Sparks, F.T., Martinez Traub, F., Daigle, T., Zeng, H., Losonczy, A., et al. (2019). Volumetric Ca(2+) Imaging in the Mouse Brain Using Hybrid Multiplexed Sculpted Light Microscopy. Cell 177, 1050–1066 e1014.

Wilt, B.A., Fitzgerald, J.E., and Schnitzer, M.J. (2013). Photon shot noise limits on optical detection of neuronal spikes and estimation of spike timing. Biophys J 104, 51–62.

Wu, J., Liang, Y., Chen, S., Hsu, C.L., Chavarha, M., Evans, S.W., Shi, D., Lin, M.Z., Tsia, K.K., and Ji, N. (2020). Kilohertz two-photon fluorescence microscopy imaging of neural activity in vivo. Nat Methods 17, 287–290.

Zhang, K., Zuo, W., Chen, Y., Meng, D., and Zhang, L. (2017). Beyond a Gaussian Denoiser: Residual Learning of Deep CNN for Image Denoising. IEEE Trans Image Process 26, 3142–3155.

Zhang, T., Hernandez, O., Chrapkiewicz, R., Shai, A., Wagner, M.J., Zhang, Y., Wu, C.H., Li, J.Z., Inoue, M., Gong, Y., et al. (2019). Kilohertz two-photon brain imaging in awake mice. Nat Methods 16, 1119–1122.

